# Patient-derived tumoroids of advanced high-grade neuroendocrine neoplasms mimic patient chemotherapy responses and guide the design of personalized combination therapies

**DOI:** 10.1101/2022.12.10.519855

**Authors:** Simon L. April-Monn, Katharina Detjen, Philipp Kirchner, Konstantin Bräutigam, Mafalda A. Trippel, Tobias Grob, Cyril Statzer, Renaud S. Maire, Attila Kollàr, Aziz Chouchane, Catarina A. Kunze, David Horst, Martin C. Sadowski, Jörg Schrader, Ilaria Marinoni, Bertram Wiedenmann, Aurel Perren

**Affiliations:** Institute of Pathology, University of Bern, 3008 Bern, Switzerland; Graduate School for Cellular and Biomedical Sciences, University of Bern, 3008 Bern, Switzerland; Charité - Universitaetsmedizin Berlin, corporate member of Freie Universitaet Berlin and Humboldt-Universitaet zu Berlin, Hepatology and Gastroenterology, Berlin, Germany; Department of Health Sciences and Technology, Eidgenoessische Technische Hochschule Zuerich, Schwerzenbach-Zuerich 8603, Switzerland; Department of Medical Oncology, Inselspital, Bern University Hospital, University of Bern, Freiburgstrasse, 3010 Bern, Switzerland; Institute of Pathology, Charité Universitaetsmedizin Berlin, Rudolf-Virchow-Haus, Berlin, Germany; Department of Medicine, University Medical Center Hamburg-Eppendorf, 20251 Hamburg, Germany; Bern Center for Precision Medicine, University & University Hospital of Bern, 3008 Bern, Switzerland

**Keywords:** Gastroenteropancreatic neuroendocrine neoplasm, NEC, Precision medicine, Drug screening, Organoid

## Abstract

There are no therapeutic predictive biomarkers or representative preclinical models for high-grade gastroenteropancreatic neuroendocrine neoplasms (GEP-NEN), a highly aggressive, fatal, and heterogeneous epithelial malignancy. We established patient-derived (PD) tumoroids from biobanked tissue samples of advanced high-grade GEP-NEN patients and applied this model for targeted rapid *ex vivo* pharmacotyping, next-generation sequencing, and perturbational profiling. We used tissue-matched PD tumoroids to profile individual patients, compared *ex vivo* drug response to patients’ clinical response to chemotherapy, and investigated treatment-induced adaptive stress responses.

PD tumoroids recapitulated biological key features of high-grade GEP-NEN and mimicked clinical response to cisplatin and temozolomide *ex vivo*. When we investigated treatment-induced adaptive stress responses in PD tumoroids in silico, we discovered and functionally validated Lysine demethylase 5A and interferon-beta, which act synergistically in combination with cisplatin. Since *ex vivo* drug response in PD tumoroids matched clinical patient responses to standard-of-care chemotherapeutics for GEP-NEN, our rapid and functional precision oncology approach could expand personalized therapeutic options for patients with advanced high-grade GEP-NEN.

## INTRODUCTION

High-grade gastroenteropancreatic neuroendocrine neoplasm (GEP-NEN), which comprise poorly differentiated neuroendocrine carcinomas (GEP-NEC) and high-grade well-differentiated neuroendocrine tumors (GEP-NET), are highly aggressive and heterogeneous cancers and there is strong need of therapies to treat them.^1–4^ Median overall survival for metastatic GEP-NECs patients is less than one year.^1–4^ Slightly better outcomes are reported in high-grade GEP-NET patients but with high and unpredictable variations in overall survival.^3^ Existing therapeutic strategies for GEP-NENs have been adopted from small-cell lung cancers (SCLC) due to their apparent clinical- and histomorphological similarities.^5–7^ Platinum-based chemotherapy is frequently used in GEP-NEC treatment. Temozolomide-based chemotherapy is currently in clinical use for high-grade GEP-NET ^8^ as response rates of platinum-based therapies seem lower.^9^

Due to the rarity and heterogeneity of the disease, extensive multi-arm clinical trials, and even exploratory and confirmatory studies, are challenging to perform. No predictive therapeutic biomarkers for high-grade GEP-NEN are in clinical use. Thus, the precise, clinical therapeutic regimes are mainly empirical, relatively uniform, and based only on small case series.^5,6^ This modus operandi has increasingly been scrutinized because uniform therapy does not account for the heterogeneity of GEP-NEN patients.^1,10,11^

Preclinical GEP-NEN models were developed during the search for predictive therapeutic biomarkers and more efficient therapy options. But these GEP-NEN preclinical models were not successfully used to develop novel or combined treatments based on mechanistic insights—this is a pressing unmet need in the field. Patient-derived (PD) xenografts of GEP-NENs have low success rates, and the few available NEN cell lines fail to accurately recapitulate the biology of high-grade GEP-NENs.^12,13^ Current GEP-NEN models provide insufficient functional and mechanistic insight into drug responses and it remains difficult to develop novel- and co-treatments for GEP-NEN patients.

We recently described a patient-derived 3-D tumoroid model that facilitates multi-center collections, efficient processing, characterization, and rapid drug screening of low abundant tumor tissues from human low-grade NET with high success rates.^14^ Sato *et al.* described a tumor organoid biobank that included stable organoid lines from a few patients with neuroendocrine neoplasms that the authors used for their longer-term cultures.^15^

Without sufficient clinical data, it is difficult to define the translational relevance of these 3-D *ex vivo* models that are derived from high-grade GEP-NEN patients. We thus set out to determine how well a patient-derived rapid *ex vivo* model recapitulates an individual patient’s response to therapy, and to test whether the rapid *ex vivo* model can provide functional insight on drug- and stress responses of individual patients.

We used targeted *ex vivo* pharmacotyping and next-generation sequencing in tumor tissues and matching patient-derived (PD) tumoroids to determine if PD tumoroids enable rapid *ex vivo* pharmacotyping and if the subsidiary biological information and the adaptive stress response patterns they provided could be used to personalize therapy strategies in individual advanced highgrade GEP-NEN patients.

We show high success rates in culturing PD tumoroids of high-grade GEP-NENs within a two-week time window. These patient-derived tumoroids recapitulated key biological features of high-grade GEP-NEN and mimicked clinical response to cisplatin and temozolomide *ex vivo.* We also investigated molecular stress responses in PD tumoroids *in silico*, discovering and functionally validating Lysine demethylase 5A (KDM5A) and interferon-beta (IFNB1)—two vulnerabilities that are synergistic in combination with cisplatin. Either KDM5A or IFNB1 can be combined with cisplatin to boost the effectiveness of the treatments, opening new therapeutic options for high-grade GEP-NENs. Together, our findings suggest we can translate patient-centered subsidiary information from PD GEP-NEN tumoroids into potentially more effective personalized treatment strategies.

## MATERIALS & METHODS

### Patient studies

We assembled a cohort of eight high-grade GEP-NEN patients from two ENETS Centers of Excellence; The University Hospital Charité Berlin (Germany) and The University Cancer Institute of the Inselspital and the University of Bern (Switzerland). Inclusion criteria were histopathologic diagnosis of G3 gastroenteropancreatic neuroendocrine neoplasm, availability of both tumor tissue- and matching cryomaterial for *ex vivo* culture, and tumor purity of >70%. A board-certified pathologist (A.P.) reviewed all cases and reclassified them according to WHO 2019 criteria (ISBN 978-92-832-4499-8) (Table 1 and Supplementary Table S1). TNM staging was based on the 8^th^ edition UICC/AJCC (ISBN: 978-1-119-26356-2). We obtained treatment and outcome information from interdisciplinary NEN tumor board records of both centers. Assessment of clinical therapy response accorded with investigator-based RECIST criteria. The final classification was based on all information about immunohistochemistry, clinical course records, and mutational status from targeted sequencing. The cohort included 3 female and 5 male patients; their ages varied from 39 to 70 years (mean = 58.0; SD = 11.8). For comprehensive patient demographics and clinical data, see Table 1, Table 2, and Supplementary Table 1. The specimens were processed as described in April-Monn, et al (2021).^14^ In brief, upon surgical resection a pathologist processed the left-over of the donor tumor tissue to 8-mm^3 cubes under sterile conditions, avoiding necrotic regions when possible. These cubes were suspended in recovery cell culture freezing medium (Thermo Fisher Scientific, USA), cryopreserved in an isopropyl alcohol freezing container (Nalgene, USA), and stored in liquid nitrogen. A consecutive block was snap-frozen (fresh_frozen) in liquid nitrogen. A mirror block was fixed in formalin and embedded in paraffin. The study was approved by cantonal authorities (Kantonale Ethikkomission Bern, Ref.-Nr. KEK-BE 105/2015) in accord with the Swiss Federal Human Research Act and by the ethics committee at Charité Universitätsmedizin Berlin (Ref.-Nr. EA1/229/17). All patients included in the study signed an institutional informed consent and agreed on the use of residual material.

**Table 1.** Patient demographics and clinicopathological classification. *At initial diagnosis, clinical differentiation between NET G3 and NEC was ambiguous. Due to an unexpected clinical course deviation, at that time a second expert opinion was obtained, which suggested a NET G3 differential diagnosis. Evaluation of the collected cryo-specimen after therapy revealed signs of acinar cell differentiation. CgA Chromogranin A MCT4 Monocarboxylate Transporter 4 PDX1 Pancreatic And Duodenal Homeobox 1 RB1 RB Transcriptional Corepressor 1 SOX9 SRY-Box Transcription Factor 9 SSTR2A Somatostatin Receptor 2 SYN Synaptophysin TP53 Tumor Protein P53 DAXX Death Domain Associated Protein ARX Aristaless Related Homeobox TRY1 Trypsin 1 BCL10 BCL10 Immune Signaling Adaptor

| Patient ID | C9502m | C8802p | C8101m | C5501m | C3301m | C0701m | aP490m | aP321m |
| --- | --- | --- | --- | --- | --- | --- | --- | --- |
| **Sex** | f | m | f | f | m | m | m | m |
| **Age at surgery [y]** | 57 | 66 | 44 | 39 | 70 | 69 | 53 | 66 |
| **Tissue source** | liver metastasis | primarius | liver metastasis | metastasis in ovary | liver metastasis | liver metastasis | lymph node metastasis | liver metastasis |
| **Primary tumor localisation** | CUP | stomach | CUP | CUP | colon | pancreas | pancreas | pancreas |
| **1-year survival** | alive | alive | alive | deceased | lost to follow-up | alive | deceased | alive |
| **Classification** | NET | NET | ACC | NEC | NEC | NET | NEC | NET |
| **Morphology subtype** | NA | NA | NA | Large-cell | Small-cell | NA | Large-cell | NA |
| **Histological differentiation** | WD | WD | NA | PD | PD | WD | PD | WD |
| **CgA** | moderate positive | NA | weak positive | strong positive | negative | moderate positive | strong positive | weak positive |
| **Ki67 [%]** | 50 | 30 | 50 | 30 | 80 | 80 | 90 | 75 |
| **MCT4** | heterogenous | NA | heterogenous | positive | heterogenous | heterogenous | heterogenous | negative |
| **PDX1** | negative | NA | negative | negative | negative | negative | negative | positive |
| **RB1** | wildtype expr. pat. | NA | wildtype expr. pat. | wildtype expr. pat. | mutant expr. pat. | wildtype expr. pat. | wildtype expr. pat. | mutant expr. pat. |
| **SOX9** | negative | NA | positiv | negative | positive | negative | strong positive | positiv |
| **SSTR2A** | moderate positive | NA | negative | weak positive | negative | negative | negative | moderate positive |
| **SYN** | strong positive | positive | negative | strong positive | strong positive | strong positive | strong positive | strong positive |
| **TP53** | mutant expr. pat. | NA | wildtype expr. pat. | wildtype expr. pat. | mutant expr. pat. | wildtype expr. pat. | wildtype expr. pat. | wildtype expr. pat. |
| **DAXX** | NA | NA | NA | NA | NA | negative | positive | negative |
| **ATRX** | NA | NA | NA | NA | NA | X | positive | positive |
| **ARX** | NA | NA | NA | NA | NA | negative | negative | positive |

**Table 2.**
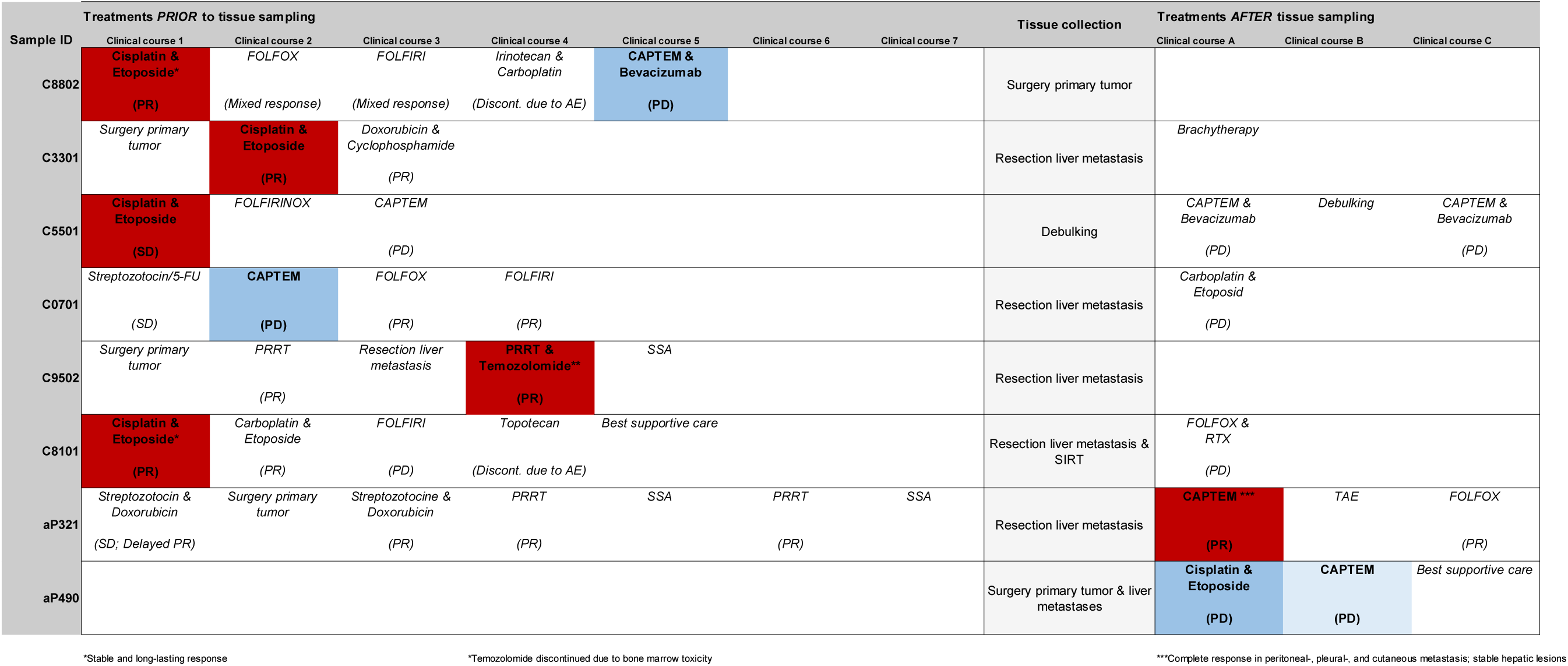
Clinical course records of GEP-NEN patients. PR partial response SD stable disease PD progressive disease

### Cancer mutation panel

The TruSight Oncology 500 Kit (TSO500, Illumina) was used for DNA library preparation and enrichment following the manufacturer’s protocol. DNA (80 ng) were sheared on a Covaris E220 ultrasonicator. DNA fragments were end-repaired, and adapters containing unique molecular identifiers (UMIs) were ligated to each fragment end. Fragments enriched by capture hybridization were analyzed by high-throughput sequencing on a NovaSeq 6000 instrument (Illumina). TSO500 alignment and variant calling was performed using the TSO500 bioinformatics pipeline v2.1.0. UMI-filtered total read counts were 103 M ± 19 M, median exon coverage was 1131 ± 253, median DNA-insert size was 136 ± 14, and % aligned reads were 98.9 ± 1.0. Sources of population frequencies that were used for auto-classification of benign variation include gnomAD (RRID:SCR_014964) and ExAC (RRID:SCR_004068). We retrieved annotations of oncogenic effects of identified variants from the OncoKB precision oncology knowledge database (RRID:SCR_014782) and assessed known activating mutations in oncogenes and inactivating mutations in tumor suppressors (Tier 1 and Tier 2). For the few cases in which IHC was inconclusive, we investigated both known mutations and mutations with unknown and unclear effects (Tier 3). OncoPrint function from ComplexHeatmap v2.6.2 (PMID 27207943) (RRID:SCR_017270) was used for visualization.

### Primary and cell line culture

For the study, we focused on naïve passage PD tumoroids to minimize clonal drift^16^ and used NEN cell line spheroids for comparison. All therapeutic studies were completed in twelve days. All screening plates contained vehicle control wells (DMSO-treated, n = 10) and blank wells (medium-only, n = 6) and for each plate, the raw fluorescent intensity values were normalized to a relative scale using the blank (B) value. Fluorescence was measured relative to the baseline of each well (BC) (Relative scale = (Fluorescence of treated cells - B)/(BC - B)).

#### Primary cell isolation and culture

Cryopreserved tumor tissues were used for *ex vivo* drug screening. For primary cell isolation, micro-cell block manufacture, and quantification, we followed the workflow described previously.^14^ In this study of high-grade GEP-NENs and in our earlier studies of lower-grade PanNENs ^14,17^, our **definition of “culture success”** for patient-derived tumoroids was based on six factors that support translational application of patient-derived GEP-NEN tumoroids: **1)** Successfully isolating and culturing viable tumor cells; **2)** retaining ± 70% of the isolated cells before drug screening; **3)** passing quality controls, including cytological, morphological, and histopathological examinations of clinically applied neuroendocrine marker expression in micro-cellblocks; **4)** attaining sufficient technical replicates (n≥4) in drug screenings; **5)** attaining stable RTG baseline and cell growth; **6)** and extending culture life spans of up to 12 days *ex vivo.*

#### NEN cell line culture

The QGP1 cell line (RRID:CVCL_3143) was purchased from the Japanese Health Sciences Foundation in 2011. QGP1 cells were kept in RPMI 1640 medium (10% FBS, 100 IU/mL penicillin, 0.1 mg/mL streptomycin). The BON1 cell line (RRID:CVCL_3985) was provided by E.J.M. Speel in 2011. BON1 cells were cultured in DMEM-F12 (10% FBS, 100 IU/mL penicillin, 0.1 mg/mL streptomycin). NT3 cells were kept in RPMI 1640 + growth factor medium (10% FBS, 100 IU/mL penicillin, 0.1 mg/mL streptomycin, 20 ng/mL EGF, 10 ng/mL bFGF) and cultured in collagen IV coated culture flasks. All cells were kept in a humidified incubator at 5% CO2 and 37 °C and cultured for no longer than two months. For all cell lines, short tandem repeat (STR) analysis by PCR was performed (QGP1 in 2011/2016/2020; BON1 in 2014/2016/2020; NT3 in 2018/2020). QGP1 cells were authenticated by their specific cancer cell profile. A BON1- or NT3-specific cancer cell profile does not exist yet, but contamination with other common cell lines can be excluded due to non-match to any known cancer cell line profile. Expression of the specific neuroendocrine markers chromogranin A and synaptophysin was routinely tested by IHC on cell blocks.

#### Compounds

Temozolomide (#S1237, Selleckchem), cisplatin (#4333164, Teva Pharma), CPI-455 (#S6389, Selleckchem), IFNB1b (#I7662-14S, Biomol) were obtained from commercial vendors and stored as stock aliquots, as indicated by the manufacturers. We selected drug concentrations for chemotherapeutics (cisplatin; temozolomide) based on physiologically relevant concentrations at each drug’s maximum tolerated serum concentration (Cmax) drawn from published human studies.^18^ We based concentrations for combinational exploratory compounds (CPI-455, IFNB1b) on primary literature and in-house *in vitro* testing of a 625-fold concentration range, optimized to induce a range of responses across classical NEN cell line spheroids (BON1, QGP1). Compounds were screened at equidistant 5-point, 625-fold concentration ranges using four technical replicates for long-term (168 hours) chemotherapeutics screens or in equidistant 3-point, 625-fold concentration ranges with three technical replicates for short-term (24 hours) combinational screens.^19^

#### *Ex vivo* drug screening

3000 - 5000 cells were plated per well. Cell viability was quantified with RealTime-Glo™ MT Cell Viability (RTG) Assay (Promega, #G9712). Assay plates were incubated for 72 hours at 37 °C in a humidified atmosphere at 5% CO2 to allow sphere formation.

#### Evaluating drug sensitivity to mono chemotherapeutics

After a baseline measurement (Day 0), we tested spheroids with titrations of cisplatin, temozolomide, or DMSO (0.16% v/v) as vehicle control. Assay plates were incubated, and RTG luminescence measurements were recorded at 96 hours and 168 hours with an Infinite 200 PRO plate reader (Tecan). A blinded experimenter scored *ex vivo* experiments and sensitivities to treatments.

#### GR metrics

Raw luminescence values were normalized to each individual baseline control value at Day 0 for the same well. *Ex vivo* responses were converted into parametrized drug sensitivity metrics, as did state-of-the-art protocols;^20^ readouts were scored on a per-division basis using the GR metrics v1.16.0 workflow described in Hafner *et al.* (2016).^21^

#### Evaluating drug sensitivity to combinational therapy

After baseline measurement (Day 0), PD tumoroids or cell line spheroids were dosed with titrations of cisplatin, IFNB1b, or CPI-455 alone or all combinations, using a full factorial drug design. We incubated assay plates and recorded luminescence measurements at 24 hours with an Infinite 200 PRO plate reader.

#### Classifying drug interaction and drug potency in combinational therapy

Our analysis was based on the combination index theorem, a mechanism-independent model for assessing drug interaction and drug potency.^19,22^ Raw luminescence values in the presence of the drug were normalized to baseline control values and DMSO-treated controls at 24 hours. Technical replicates were averaged to yield a mean relative cell count per condition. From this, we calculated observed inhibition (%) and fractional inhibition effects (fa) for visualization and downstream analysis and then used the CompuSyn v1.0 workflow to extract drug interaction and drug potency metrics.^22^ We summarized the degree of drug interaction by drug combination indices (CI), and then used an isobologram to describe how the drug combination activity we observed deviated from isoactive monotherapies.^19^ The median-effect equation was used to derive drug potency parameters (Dose Reduction Index [DRI]; Effect at Dose X [DE]).^19^

### Nucleic acid extraction

A DNA Purification Micro Kit (Norgen Biotek, #50300) was used to extract genomic DNA from fresh frozen tumor tissue. Total RNA was extracted from fresh frozen tumor tissue or cultured cells with a Single Cell RNA Purification Kit (Norgen Biotek, #51800). Nucleic acid quantification was performed with the Qubit DNA/RNA HS detection kit (Thermo Fisher Scientific, #Q32852). We used a Femto Pulse system with an Ultra Sensitivity RNA kit (Agilent, #FP-1201-0275) to analyze quality control metrics.

### Immunohistochemistry

All the immunohistochemistry (IHC) markers were repeated on freshly cut tissue blocks and re-evaluated by a NEN expert pathologist (A.P.). For immunohistochemistry, we cut the paraffin-embedded material into 2.5-μm-thick serial sections and then deparaffinized, rehydrated, and retrieved antigens with an automated immunostainer (Bond RX, Leica Biosystems). Antigen retrieval was performed in a Tris-EDTA buffer for 30 min at 95°C for Ki-67 (1:200, Dako, M7240), ATRX (1:400, Sigma-Aldrich, HPA001906), MCT4 (1:50, Santa Cruz, sc376140 D1), SOX9 (1:100, Cell Signalling, 82630T D8G8H), ARX (1:1500, R&D Systems, AF7068), PDX1 (1:100, R&D Systems, MAB2419); in a Tris-EDTA buffer for 30min at 100°C for synaptophysin (1:100, Novocastra, 27G12), CgA (1:400, CellMarque, 238M-94 LK2H10), SSTR2A (1;50, BioTrend, SS-8000-RM UMB-1); in a proteinase K solution for Trypsin 1 (1:20000, Chemicon, MAB1482); in a citrate buffer for 30min at 100° for DAXX (1:40, Sigma-Aldrich, HPA008736), RB1 (1:200, BP Pharmingen, 554136 G3-245); and, in a citrate buffer for 20 min at 95°C for TP53 (1:800, Dako, M7001 DO-7), BCL-10 (1:1000, Santa Cruz, sc-5273 331.3). Primary antibody incubation lasted 30 min at the specified dilutions. Visualization used a Bond Polymer Refine Detection Kit (Leica, #DS9800) (RRID:AB_2891238) for visualization; DAB (3,3’-Diaminobenzidine) was the chromogen. Slides were counterstained with hematoxylin. We used an automated slide scanner Panoramic 250 (3DHistech) at 20x magnification to capture scans and acquired images with QuPath software (PMID: 29203879).

### Bulk RNA sequencing

#### Library preparation and sequencing

Sequencing libraries were prepared from RNA using the SMARTer Stranded Total RNA-Seq Kit v3 for picogram input material (Takara, #634488). Libraries were sequenced as paired-end 101 bp (tumoroid samples) or paired-end 81 bp (original tumor tissues) reads on a NovaSeq 6000 (Illumina) platform at ~30M reads/sample. Reads were demultiplexed and converted to FASTQ format with bcl2fastq v2.20.0.422 (RRID:SCR_015058). Cutadapt v2.5 (PMID 28715235) (RRID:SCR_011841) was used to trim Illumina adapter sequences and mask 3’ homopolymers longer than 10 bp. We removed reads containing more than 20 masked bases or shorter than 65 bp (tumoroid samples) or 50 bp (original tumor tissue). Trimmed reads were mapped against a custom list of ribosomal RNAs and repetitive RNA elements with bwa v0.7.17 (PMID 19451168) (RRID:SCR_010910); mapping reads to this custom list were discarded. At each step, we used FastQC v0.11.7 (RRID:SCR_014583) to track read quality. Processed reads were mapped to the human genome (GRCh37, GENCODE annotation v37) with STAR v2.7.3a (PMID 23104886) (RRID:SCR_004463). Mapped reads were deduplicated based on the 8bp UMI in the R2; we used UMI-tools v0.5 (PMID 28100584) (RRID:SCR_017048) and the default directional method. Deduplicated reads were assigned to GENCODE v37 genes in subread v2.0.1 (PMID 24227677) (RRID:SCR_009803). We excluded one drug-treated sample from the tumoroid culture of patient C5501m because input and library quality was low.

#### Differential gene expression

To compare original tumor tissue and tumoroids, we normalized expression data with smooth quantile normalization (qsmooth v1.8.0) (PMID 29036413). To determine differentially expressed genes, we combined limma voom (PMID 25605792) (RRID:SCR_010943) with the duplicateCorrelation function to model repeated measurements of the same patient. For drug-treated tumoroids, we determined differential expression with DESeq2 v1.32.0 (PMID 25516281) (RRID:SCR_000154). Earlier gene expression studies found that focusing on sublethal drug concentrations prevented artificially exaggerating non-specific cellular stress or death processes caused by high drug dosages.^23,24^ Hence we analyzed sublethal concentrations of cisplatin (0.53uM) and temozolomide (11.52uM) to determine the drug-related mode of action. Treatment-independent expression variability was modeled using surrogate variable analysis (SVA) from sva v3.40.0 ^25^ (PMID 17907809) (RRID:SCR_002155). All available surrogate variables were added to the DESeq2 model. Log2 expression fold changes of highly variable genes were shrunk with the apeglm v1.14.0 algorithm (PMID 30395178). Differentially expressed genes from cisplatin treatment were analyzed in metascape (PMID 30944313).

#### Hierarchical clustering and consensus clustering

We used the consensus clustering algorithm to investigate the transcriptional similarity between the original tumor and patient-derived tumoroids. Original tumor tissues and tumoroids were consensus clustered with ConsensusClusterPlus v1.54.0 (PMID 20427518) (RRID:SCR_016954) on the Pearson correlation of the 2000 most variable genes (innerLinkage: Ward.D2, finalLinkage: Average).

#### Master regulator protein activity analysis

Master Regulator (MR) proteins represent mechanistic determinants of a tumor’s transcriptional states.^26^ We leveraged a pan-GEP-NEN regulatory network (context-specific interactome) from transcriptional profiles of GEP-NEN patient samples (n=212) (GSE98894). We used ARACNe (accurate reconstruction of cellular networks) (PMID 15778709) (RRID:SCR_002180) to reverse-engineer regulatory networks and VIPER (Virtual Proteomic by Enriched Regulon analysis)^26^ to transform transcriptional profiles into master regulators protein activity profiles and to infer master regulator protein activity in original tumors and patient-derived tumoroids (n=8) from our GEP-NEN patients.

#### Functional enrichment analysis

UpSetR v1.4.0 (PMID: 28645171) was used to visualize intersections between gene sets. To compare original tumor tissue and tumoroids, we selected differentially expressed genes (adjusted p-value < 0.05), and tested enrichment of Gene Ontology terms in topGo v2.44.0 (PMID: 16606683) (RRID:SCR_014798) (Kolmogorov-Smirnov, adjusted p-value < 0.01). The GO graph was modeled by the weight01 algorithm; terms with less than five members were excluded. Gene set enrichment analysis (GSEA) (RRID:SCR_003199) was performed in clusterProfiler v.3.18.1 (PMID: 22455463) (RRID:SCR_016884) based on log2 expression fold changes.

#### Perturbational profiling in cMap

We compared the top and bottom 150 genes from drug versus control-treated tumoroids (adjusted p-value < 0.05, sorted by the Wald statistic) to the compendium of perturbational reference signatures from Connectivity Map (L1000, Touchstone v1.0)^27^ (RRID:SCR_015674) and extracted connectivity map scores (τ) for all available knock-down (kd), overexpression (oe), and compound perturbagens. To estimate the robustness of matching signatures, we rarified or permutated the input lists of differentially expressed genes.

### Data availability

Sequence data that support the findings of this study have been deposited in Gene Expression Omnibus (GEO); primary accession code is GSE213504. We adapted the code supporting our study from published R packages and other software. All code is available from the corresponding author on request.

## RESULTS

### In-depth characterization of the high-grade GEP-NEN patient cohort

To investigate whether patient-derived tumoroids can successfully model advanced malignant GEP-NENs and to elucidate the biology of the disease, we conducted a retrospective cohort study of human high-grade GEP-NEN patients who were operated at the University Hospital of Bern (CH) or Charité University Hospital Berlin (DE). During a systematic retrospective review of hospital biobank records, we identified and retrieved those cases in which fresh-frozen tissue-matched cryopreserved G3 NEN tumor tissues were available. Between 1987 to 2022, we identified eight patient cases from the cryopreserved GEP-NEN patient samples (n=311) (Fig. 1A). This small sample reflects the rarity of this type of tissue resource, as most patients are diagnosed at an advanced metastatic stage and undergo diagnostic biopsies rather than surgery.

**FIGURE 1.**
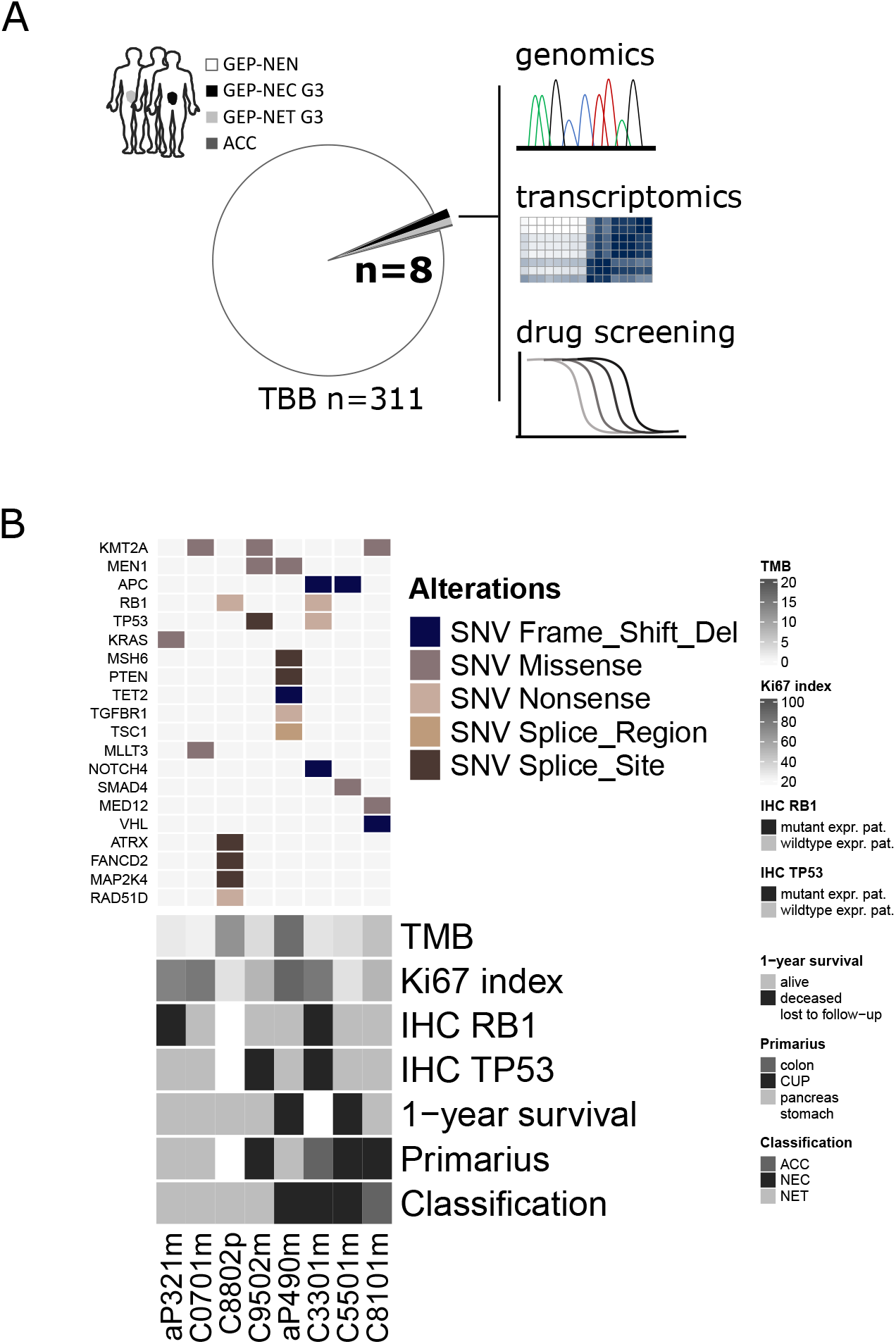
Study overview and clinical presentation of high-grade GEP-NEN patient cohort. **A)** Schematic diagram of study outline, material processing, and analysis performed in the present study. TBB tumor biobank **B)** Oncoplot showing common genetic alterations of GEP-NENs found in the study cohort together with a selection of clinical parameters. The upper panel indicates specific types of single nucleotide variations (SNV) found in fresh frozen original tumor tissue from high-grade GEP-NEN patients. The lower panel displays the patients’ clinical parameters, including tumor mutational burden (TMB; mutation/Mb) and 1-year survival, proliferative activity (Ki-67; percentage positive cells per tissue), RB1 protein expression, TP53 protein expression, location of primary tumor, and the diagnostic classification. NET neuroendocrine tumor, NEC neuroendocrine carcinoma, ACC acinar cell carcinoma CUP cancer of unknown primary Mutant expr. pat. Mutant expression pattern (TP53 loss of protein (0% positive tumor cells) or overexpression (≥90% positive tumor cell; RB1 complete loss of protein), wildtype expr. pat. wildtype expression pattern.

The cohort comprises high-grade metastatic neuroendocrine tumors (NET G3, n=4), neuroendocrine carcinomas (NEC, n=3) of gastric-, pancreatic- (Pan), or unknown primary (CUP) site, and one additional case that had been diagnosed as NEN (n=1) at the time of initial diagnosis, but the liver metastasis that we obtained at a later disease stage was reclassified as acinar cell carcinoma during our case review (Table 1). Patient demographics, clinicopathological classification, and comprehensive clinical course records are presented in Tables 1 and 2 and Supplementary Table S1.

The patients’ fresh frozen material was subjected to a transcriptomic molecular analysis and nextgeneration sequencing to profile the tumor’s cancer-related gene mutational burden (Supplementary Fig. S1A). The tumor mutational burden (TMB) in all patients was low (3.1 mt/Mb; IQR 5.6 mt/Mb; median; IQR) except in two patients whose TMB was elevated (aP90m 11.8 mt/Mb; C8802p 16.4 mt/Mb) (Supplementary Table S1). Microsatellite instability (MSI) was low (2.4% ± 1.9%; mean ± SD), and we detected no alterations in copy number (Supplementary Table S1). The most frequent single nucleotide variants (SNV) were missense mutations. Among the SNVs in our samples, we found well-known prototypic genetic drivers of GEP-NET (MEN1, ATRX) and GEP-NECs (TP53, RB1, APC, SMAD4) (Fig. 1B, Supplementary Table S2), supporting histopathological diagnosis (Table 1 and Supplementary Fig. S1A).

### Phenotypic characteristics of high-grade GEP-NEN patient-derived tumoroids resemble original tumor tissue

We successfully generated PD tumoroids from all cryopreserved tissue-matched specimens based on the criteria specified in our methods section and found support for translational application of GEP-NEN PD tumoroids (*see Methods*, page 7). We first determined if PD tumoroids preserve relevant histomorphological features of original high-grade GEP-NENs in culture. Two board-certified pathologists (A.P., M.T.) confirmed PD tumoroids were alike the original tumor tissue in high tumor content, in tumor cell morphology, and in the expression of diagnostic neuroendocrine biomarker synaptophysin, based on the cytology of micro-cell-blocks from cultured cells (Fig. 2A, Supplementary Fig. S2A, Supplementary Table S3). Thus, patient-derived tumoroids did preserve the neuroendocrine phenotype of GEP-NEN tumor cells. Moreover, we detected a focal presence of extracellular matrix (C9502m, C8802p, C5501m) and calcifications (C9502m) (Supplementary Fig. S2B, Supplementary Table S3). Since we intentionally depleted stromal cells in the 3-D culture workflow, non-neoplastic cells - including fibroblasts and macrophages - were less abundant in PD tumoroids than in original tumor tissue (Supplementary Fig. S2B, Supplementary Table S3). Patient-derived tumoroids also exhibited increased metabolic activity *ex vivo* over time (Supplementary Fig S2C), and this increase significantly correlated with the proliferation indices in the donor tissues and the patients’ clinical tumor grades (Supplementary Fig. S2D).

**FIGURE 2.**
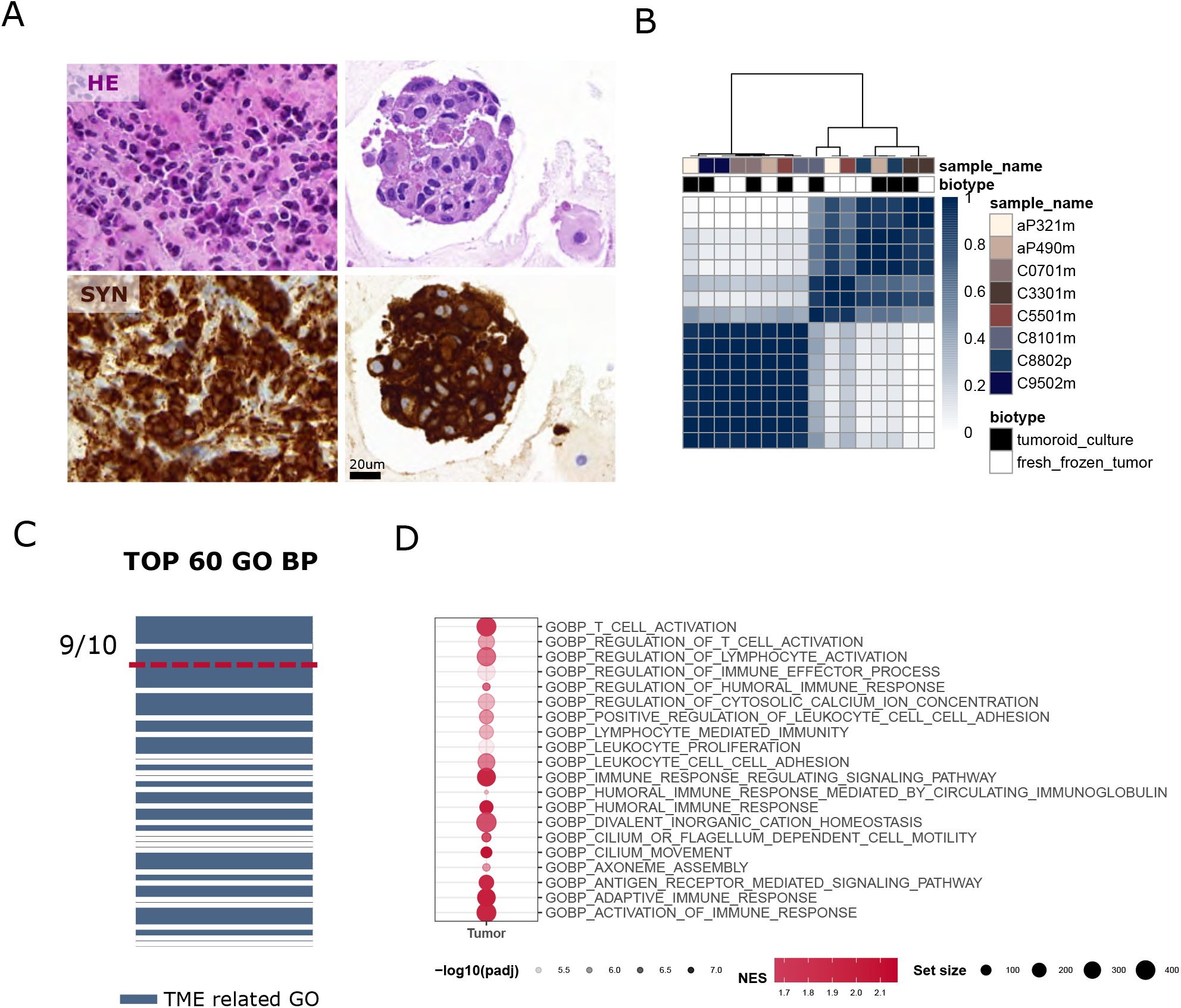
Patient-derived tumoroids recapitulate biological key features of original tumors. **A)** Representative Hematoxylin and eosin (HE) staining and neuroendocrine marker synaptophysin (SYN) immunolabeling in original tumor tissue and tissue-matched patient-derived (PD) tumoroids. Scale bar in all stainings, 20 μm. **B)** Consensus clustering of original tumor tissue and PD tumoroids according to the 2000 most variable genes by variance based on RNA sequencing. Cluster stability was reached for k = 2. Consensus cluster correlation is represented by the blue scale. Each column represents one sample. Biotypes and patient identifiers are colored by class. The heatmap inner linkage was determined by Ward.D2 and the dendrogram (outer linkage) by an average of correlation scores. **C)** Gene ontology (GO) enrichment comparing original tumor tissues and PD tumoroids. Displayed are the 60 most significantly enriched GO terms of biological processes (p-ks < 0.005, Kolmogorov-Smirnov test). Tumor-micro-environment (TME) related GO terms are indicated in blue. In the top ranks (red line), nine GO terms are directly linked to the immune cell compartment. **D)** Gene set enrichment analysis comparing original tumor tissues with PD tumoroids. Displayed are the top-ranked and most significantly enriched gene sets (GO biological processes) found in original tumor tissue. Dots represent GO term enrichment: Red color indicates normalized positive enrichment score (NES) > 1.7; Transparency indicates Benjamini-Hochberg adjusted p-values (p-adj); Size indicates the number of genes within the specific gene set.

We used next-generation RNA sequencing to assess the extent to which transcriptional expression patterns of original tumors had been retained in matching PD tumoroids. The consensus among gene expression of original tumor tissues and tissue-matched PD tumoroid pairs was high, as evidenced by an unsupervised consensus cluster analysis of the top 2000 most variable genes (Fig. 2B). Additionally, inferred protein activity signatures of master regulators were largely conserved in original tumor tissue and patient-derived tumoroid pairs and reconfirmed a high similarity between both biotypes (Supplementary Fig. S3A).

Principal component (PC) analysis of original tumor tissue and PD tumoroids made clear that patient-specific expression patterns had been systematically retained in culture (Supplementary Fig. S3B). Also commonly occurring cancer hallmark pathways (Myc Targets V1, G2M checkpoint; Oxidative phosphorylation; Protein secretion) as well as pathways related to P53 regulation or EGF- and VEGF signaling were retained based on the gene expression analysis (Supplementary Fig. S3C). In contrast, gene expression in spheroids grown from two conventional NEN cell lines (QGP1 and NT3) markedly diverged from patient samples (Supplementary Fig. S3D).

Interestingly, PC1 (21% of total variance) separated PD tumoroids from original tumor tissues (Supplementary Fig. S3B). To explore this further, we investigated differentially expressed genes in original tumor tissues and matched PD tumoroids using functional enrichment analysis. The largest fraction of significant biological process gene ontology (GO) terms was linked to the tumor microenvironment, especially the immune cell compartment (9/10 Top10 GO BP) (pks < 0.005) (Fig. 2C; Supplementary Fig. S3E; Supplementary Table S4), which reconfirmed the cytology findings that there were very few immune- and/or stromal cells in our patient-derived GEP-NEN tumoroids (Supplementary Fig. S2B, Supplementary Table S3). In the pathways most enriched in original tumor tissue (NES > 1.7, p-adj < 0.005), adaptive- and innate immunity-, interleukin- and cytokine-related GO terms were strongly overrepresented (59/91 pathways), indicating that the depleted immune cell compartment accounts for much of this difference (Fig. 2D, Supplementary Table S4).

Altogether, transcriptomic profiles, histomorphology, and functional readouts underlined that GEP-NEN PD tumoroids are biologically complex and retain key traits of original GEP-NEN donor tumors, and that PD tumoroids harbor a degree of histological, cellular, and molecular diversity closer to original tumors than conventional permanent NEN cell lines.

### High-grade GEP-NEN patient-derived tumoroids mimic clinical response to platin and temozolomide treatment *ex vivo*

To test if drug sensitivities in PD tumoroids mimicked clinical patient responses, we performed *ex vivo* drug pharmacotyping in all samples. Based on established first-line therapy recommendations for GEP-NEC and high-grade GEP-NET patients,^5^ we screened all PD tumoroids for their *ex vivo* sensitivity to cisplatin (CPT) or temozolomide (TEM) chemotherapy (Fig. 3A, Supplementary Table S5). Then we converted *ex vivo* responses from naïve-passage PD tumoroids into parametrized drug sensitivities using GR metrics ^21^ to account for differences in proliferation rates among samples, based on state-of-the-art protocols for tumor organoid and other 3D-culture screens.^20^

**FIGURE 3.**
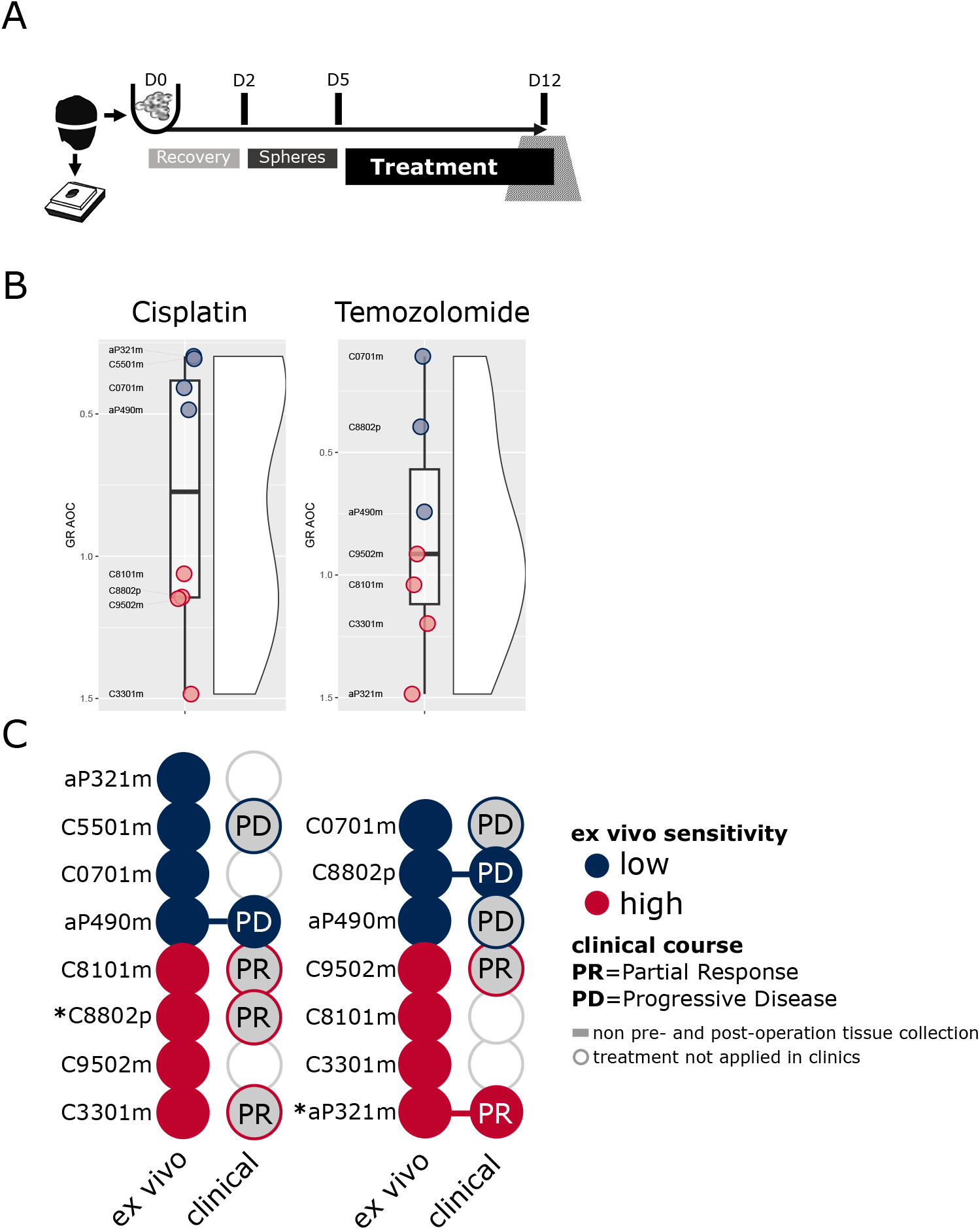
Patient-derived GEP-NEN tumoroids mimic the clinical patient response. **A)** Schematic diagram of *ex vivo* drug screening workflow in patient-derived GEP-NEN tumoroids. D day(s) **B)** Effect of cisplatin and temozolomide treatment on viability in PD tumoroids. PD tumoroids were treated with DMSO (Control), cisplatin, or temozolomide for 168 hours. *Ex vivo* drug sensitivities were converted into parametrized drug sensitivity metrics using GR metrics (GR AOC). **C)** Comparison between *ex vivo* sensitivity of PD tumoroids and clinical patient response for cisplatin (left) and temozolomide (right). Circles connected with lines represent patients with clinical therapy results adjacent to pre-/post-operative specimen collection. * These two patients (C8802p and aP321m) clinically showed accentuated and long-lasting responses to either cisplatin or temozolomide systemic therapy.

For both treatments, PD tumoroid drug sensitivities varied between patients (Supplementary Fig. S4A+B), showing a spectrum of drug sensitivity (Fig. 3B+C). The *ex vivo* sensitivity we observed in PD tumoroids was consistent with the patient’s response to clinical therapy (Table 2). For those cases in which we could directly compare the patient’s nearest clinical responses - (± 2 months) post- and/or pre-operative to the cryo-specimen collected from the patients - we found sensitivity in the PD tumoroids mimicked clinical patient responses for both temozolomide (n=2) and cisplatin (n=1) therapy (Fig 3B+C). The functional readout derived from the screen also complemented the pathological and clinical features (Ki-67 index, differentiation, TP53/RB1/KRAS mutational status, MGMT promoter methylation status) (Supplementary Fig. S4B) that recommendations suggest be consolidated *before* selecting a therapy in individual cases.^5^

Patients whose response to systemic therapy was accentuated and long-lasting (C8802p and aP321m) also exhibited high *ex vivo* drug sensitivity (Fig. 3B+C, Table 2). PD tumoroids from these patients were exclusively sensitive to either cisplatin- or temozolomide-based treatment but not both (Fig. 3B+C), which aligned with their clinical records. These findings suggest that patient-specific drug sensitivities and inter-patient susceptibilities are retained in PD GEP-NEN tumoroids and that cultured PD GEP-NEN tumoroids provide sensitive and direct functional information on *ex vivo* drug responses in individual patients.

### Transcriptional perturbational profiling in high-grade GEP-NEN PD tumoroids defines adaptive stress response to chemotherapy

We then sought to determine if molecular perturbation profiles from PD tumoroids could provide mechanistic insights into adaptive stress responses and reveal novel treatment vulnerabilities. To accomplish this, we generated transcriptional perturbation profiles from matched PD tumoroids after DMSO control, cisplatin, or temozolomide treatment. PCA of these profiles revealed that patient-specific expression differences were greater than cisplatin- or temozolomide-induced expressional effects (Supplementary Fig. S5A). Grouping the cohort based on changes in their global gene expression did not clearly separate PD tumoroids with high- and low cisplatin- or temozolomide *ex vivo* sensitivities. (Supplementary Fig. S5B). Neither did the magnitude of gene expression changes correlate with *ex vivo* sensitivity (Supplementary Fig. S5C). We considered that treatment-independent sources of variation precluded the detection of correlations and hence sought to factor in such sources using surrogate variable analysis (SVA).^25^ As we expected, these surrogate variables correlated with known biological variables, including patient age, gender, tumor type, Ki-67 index, and sequencing depth (Supplementary Fig. S4D+E).

Notably, we found *ex vivo* sensitivity was associated with the surrogate variables (Supplementary Fig. S5D); testing differential gene expression and factoring in all surrogate variables, yielded a clear cisplatin-induced perturbation signature (327 DEGs, FDR = 0.1, p-adj < 0.05) (Fig. 4A), and greatly enriched significant p-values in the p-value distribution (Supplementary Fig. S4F). Since differential expression results for temozolomide were smaller (28 DEGs, FDR = 0.1, p-adj < 0.05, Supplementary Fig. S5F), we focused only on cisplatin in our subsequent analyses.

**FIGURE 4.**
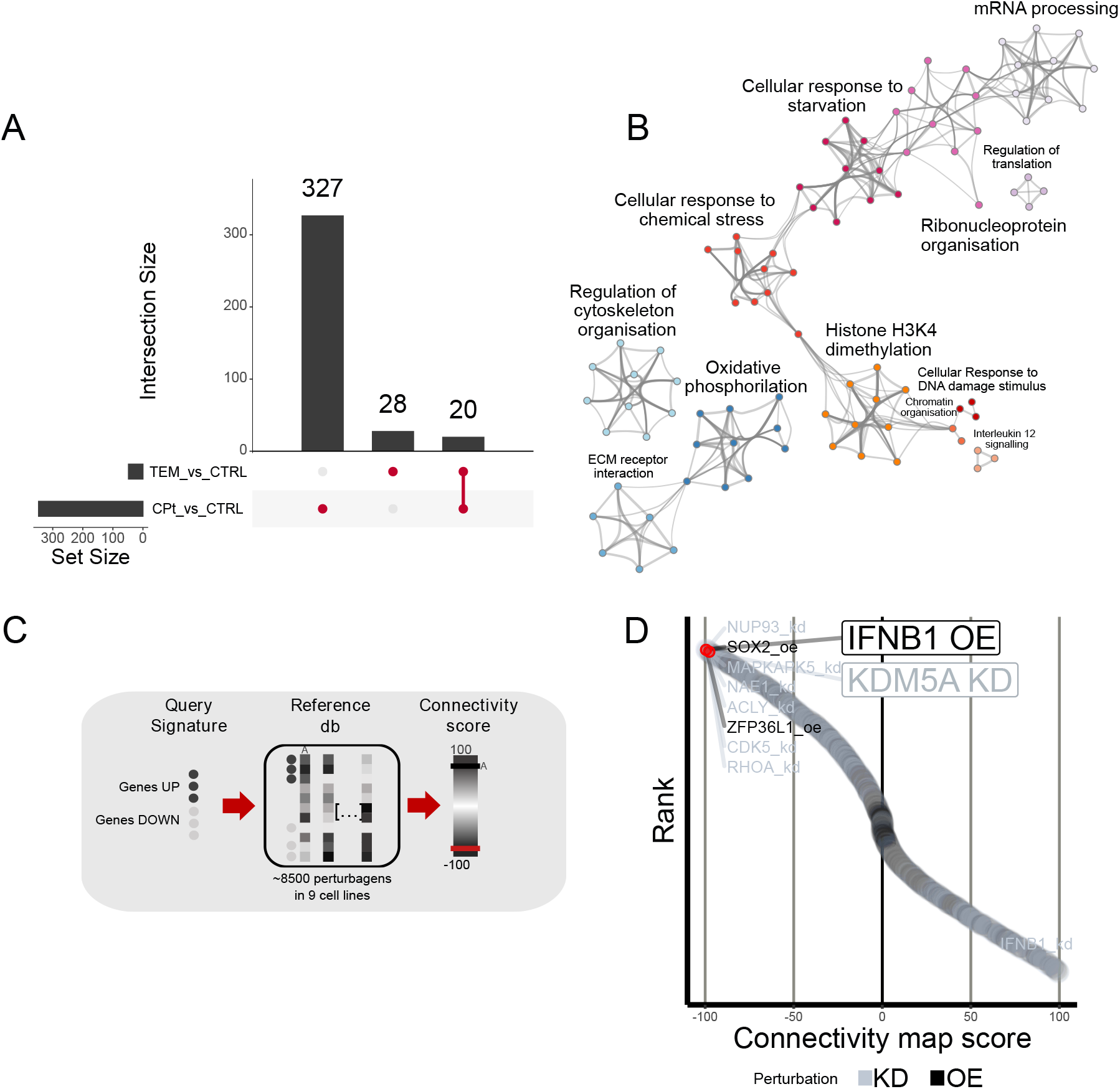
Molecular stress response in patient-derived tumoroids reveals IFNB1 and KDM5A as targets for combination therapy with cisplatin. **A)** UpSet plot of differentially expressed genes in patient-derived (PD) tumoroids treated with cisplatin (CPT) or temozolomide (TEM). Highlighted are the number of genes specific for either CPT or TEM stress response or genes shared between both. **B)** Network representation of pathway and GO biological processes from cisplatin-induced stress response genes (n=327). Each node represents an enriched term and is colored by its cluster. **C)** Schematic diagram of connectivity map (cMap) workflow to detect connectivity between stress response signatures from PD tumoroids (top 150 up- or down-regulated genes) and perturbational signatures in the database. **D)** Waterfall plot of cMap targets inducing inverse response signatures (connectivity map score (τ) < 0) to cisplatin-induced stress response in PD tumoroids. The list of top hits (red rectangle) is magnified (right). τ stands as a standardized measure ranging from −100 to +100; A τ of −90 indicates that only 10% of reference perturbations showed stronger connectivity to the query ^27^. Overexpression (oe) of IFNB1 and knockdown (kd) of KDM5A induce highly inverse signatures to the cisplatin-induced stress response.

Pathway (REACTOME; KEGG; WIKI) and GO gene set enrichment analysis on cisplatin-induced perturbation signatures revealed well-known underlying biological themes such as response to chemical stress or DNA damage (Fig. 4B), DNA repair, and apoptosis (Supplementary Fig. S5G).^28^ Histone H3K4 methylation also prominently contributed to the perturbation gene signature (Fig. 4B), suggesting the possible relevance of epigenetic targets. To explore the underlying biological themes further, we compared the cisplatin-induced perturbational signature to the Connectivity Map (cMap) ^27^, a large perturbation signature database (Fig. 4C). When we focused on pathways annotated in cMap as “DNA directed compounds,” we found Amonafide (a DNA intercalating agent) was among the top-ranked compounds and had very high connectivity score (τ = 96.05) while temozolomide (a DNA alkylating agent) had a nearly neutral connectivity score (τ = −6.38). These findings corroborate the specificity of the cisplatin-induced perturbation signature (Supplementary Fig. S5H, Supplementary Table S6).

### IFNB1 and KDM5A genetic perturbation induces inverse expression signatures to cisplatin chemotherapy of high-grade GEP-NEN PD tumoroids

Cancer escape mechanisms and the inevitable emergence of resistance to monotherapies make it imperative to formulate effective combinational chemotherapies, which are now fundamental to modern cancer therapy. ^29–33^ We used transcriptional perturbation profiles from treated patient-derived GEP-NEN tumoroids to infer targets for combination and examine the cisplatin-induced perturbation signature to identify possible combinational treatment options. To prioritize and evaluate complementary combinations, we examined perturbation candidates that created gene expression signatures inversely related to cisplatin-treated PD tumoroids. Overexpression of Interferon Beta 1 *(IFNB1)* and knock-down of Lysine Demethylase 5A *(KDM5A)* in cMap’s core cell panel (3147 genetic perturbations) were among the top-ranked perturbational candidates, with highly inverse connectivity map scores *(IFNB1*, rank 15, τ = −99.54; *KDM5A*, rank 52, τ = −97.68) (Fig 4C; Supplementary Table S6) and this pattern was robust and specific (Supplementary Fig. S5I; Supplementary Table S6). Both, KDM5 isoforms and IFNB1 receptors (IFNAR1 and 2) mRNA, were expressed in PD tumoroids (Supplementary Fig. S5J). Together, these findings indicate that molecular stress responses in PD tumoroids are specific and can be exploited to, *in silico*, predict treatment vulnerabilities.

### *In silico*-predicted combinational therapies induce effective and synergistic treatment responses in patient-derived GEP-NEN tumoroids

To evaluate the functional activity of *in silico*-predicted candidates in combinational drug therapy, we applied either human recombinant IFNB1 or KDM5A-inhibitor with cisplatin in high-grade PD tumoroids and NEN cell line spheroids. We used the combination index theorem to analyze synergistic drug interaction and combined drug potency.^19,22^ The degree of drug interaction was summarized as a drug combination index (CI) and the drug potency was based on the inhibitory effect and the dose reduction achieved in the combination treatment (Fig. 5A). We found that highgrade GEP-NEN tumoroids were susceptible to mono- and combinational treatment with recombinant INFB1 or KDM5A inhibitor (Supplementary Fig. S6A+B). We then used the overall inhibitory effect of cisplatin monotherapy at a physiologically relevant concentration (Cmax 14.4 uM; inhibition 0.29 ± 0.24, mean ± SD) to select a reference level for comparing drug interaction and drug potency among tumoroids.

**FIGURE 5.**
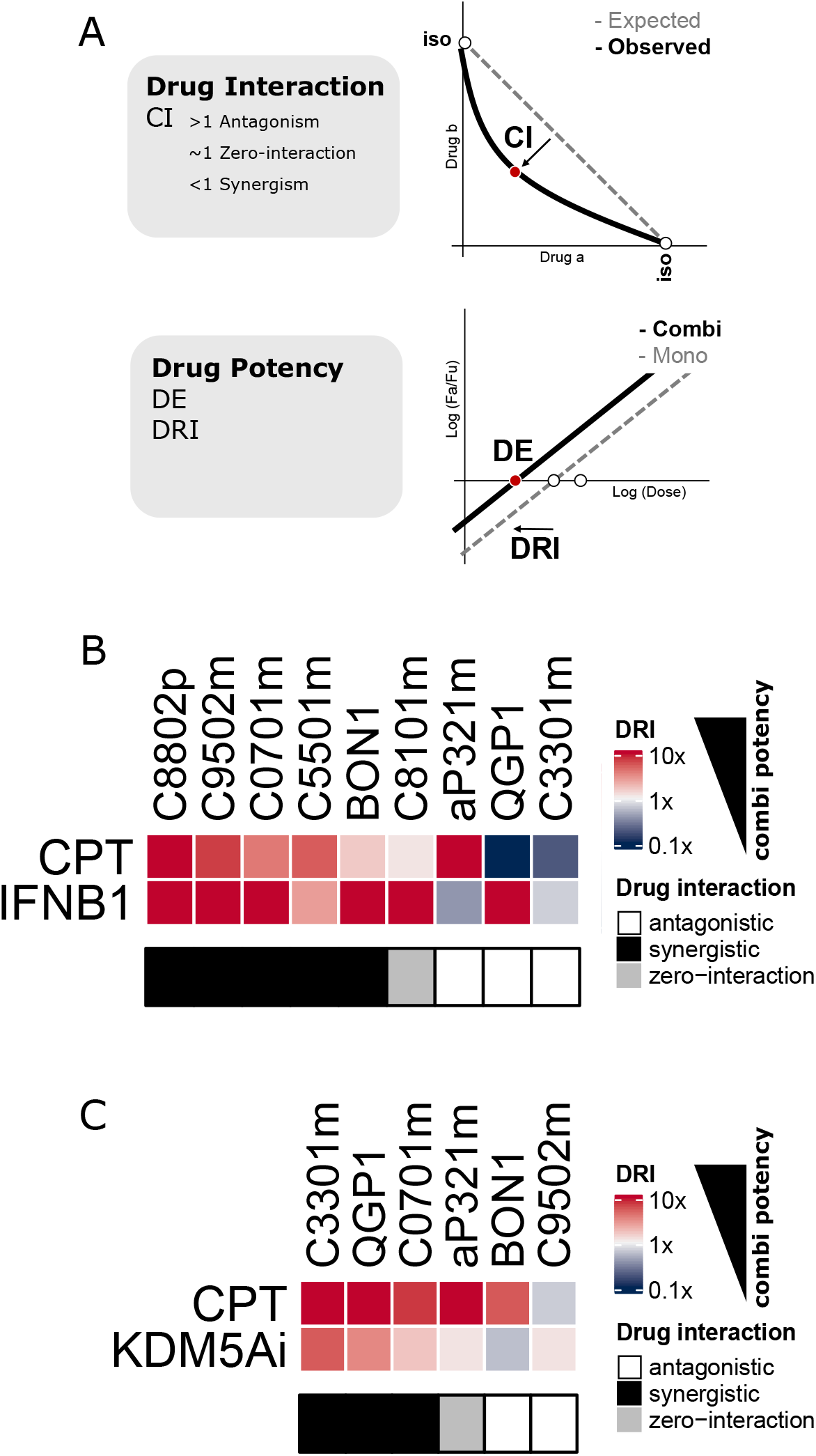
Combinational treatment of cisplatin and KDM5A or IFNB1 induces synergistic and potent treatment response *ex vivo*. **A)** Schematic representation of parameters to assess combinational therapy. Drug interaction was assessed by determining the combinational index (CI) as the deviation of the observed drug combination activity from isoactive monotherapies (iso) at a defined effect level. **Drug potency** parameters were derived from the median-effect equation by calculating the dosage required to reach a specific effect level (DE) and by calculating the dose reduction index (DRI), describing the fold-change of required dosage in combination therapy compared with the required dosage in respective monotherapy. **B+C)** Heatmap displaying drug potency parameters for combinational therapy in patient-derived (PD) tumoroids. The color scale indicates the drug reduction index (DRI) relative to the combination treatment. Red (high DRI) or blue (low DRI) indicate higher or lower drug doses required for an equipotent inhibitory effect in monotherapy. Drug interaction was classified into synergistic (black), antagonistic (white), or zero-interaction (grey) based on CI.

In line with our *in silico* findings, exposure to CPT+INFB1 combination indicated synergistic drug interaction in five of the screened tumoroids *ex vivo* (5/9) (CI=0.43 ± 0.32, mean ± SD) (Supplementary Fig. S6C). Similarly, exposure to the CPT+KDM5A combination yielded synergistic drug interaction in three tumoroids *ex vivo* (3/6) (CI= 0.43 ± 0.23, mean ± SD) (Supplementary Fig. S6D). In PD tumoroids where synergy was detected, combinational dosages needed to obtain equipotent inhibitory effects were considerably lower than in monotherapies (Supplementary Fig. S6E+F), showing highly favorable dose-reduction indices (DRI >> 1) for each individual drug (Fig. 5B+C) and emphasizing the increased potency of combination therapy *ex vivo.*

Altogether, our findings show that NEN PD tumoroid *ex vivo* drug screening and perturbational profiling can be successfully applied for the timely assessment of standard-of-care therapies and the likely effects of experimental drugs. Our analysis of therapy-induced adaptive stress responses revealed two clinically attractive co-vulnerabilities, which proved that our findings have direct functional significance for patient-derived and cell line GEP-NEN tumoroids.

## DISCUSSION

Therapeutic target discovery, validation, and translational applications face severe obstacles in rare cancers such as high-grade GEP-NEN. The selection of therapies for high-grade GEP-NENs is largely based on clinical experience in the absence of large clinical trials ^1,11^ and the absence of predictive biomarkers for therapy.^5^ Our data demonstrate that high-grade GEP-NEN PD tumoroids are well suited for rapid *ex vivo* pharmacotyping and provide biological information on this lethal malignancy. Pharmacotyping may also provide useful therapeutic information, helping oncologists select the best therapies for high-grade GEP-NENs, even in the absence of clinical trials^1,11^ and predictive biomarkers.^5^

The lack of existing preclinical disease models is a major obstacle in the study of rare cancers. ^12,13^ For the first time, we combined a patient-derived model system of high-grade GEP-NEN, extensive characterization of matched tumor tissues, and comprehensive patient clinical follow-up for the study of these rare cancers. Cytology confirmed that our PD tumoroid cultures had high tumor content intermixed with few non-neoplastic cells, including fibroblasts and macrophages. The functionality of the PD tumoroids was evident in the retention of neuroendocrine protein expression and high *ex vivo* proliferation (Fig. S2). Hence, PD tumoroids provided a faithful representation of these defining features of high-grade NENs. Inter-patient molecular transcriptional patterns were retained in tissue-matched PD tumoroids, further demonstrating that key biological features are recapitulated in the model (Fig. 2). The relevance of patient-derived models is underscored by the clear difference between transcriptomes of classical NEN cell line compared to the patient material and PD tumoroids.

Providing clinicians with evidence obtained from patient-specific models may help them offer personalized treatment. PD tumoroids of high-grade GEP-NEN patients mimic patient response to established first-line chemotherapies (Fig. 3, Supplementary Fig. S4). While our cohort was small, our results align with those of similarly sized studies of similar sizes of various cancer entities that demonstrated clinical applicability using patient-derived *ex vivo* models, e.g., in colorectal cancer,^34,35^ pancreatic cancer,^36^ and lung cancer.^37^

We efficiently and successfully processed low abundant GEP-NEN tissues with minimal cell requirements, included critical quality control steps, and ensured turnaround time was only two weeks (Fig. 2+3, Supplementary Fig. S2-4)—far less than the 2 to 6 months reported in other precision medicine studies.^37,38^ Because patients with high-grade NEN are not expected to live long without effective treatment, a rapid turnaround using PD tumoroids better matches the clinical course of these patients. Especially in high-grade NENs, when survival times are expected to be short, fast turnaround may facilitate timely therapy decisions that can extend patients’ lives. As a trade-off for rapid information, our workflow and model are focused on “one round of experimentation” per preparation. The rapid *ex vivo* culture and analysis of individual tumor specimens can however be expanded with additional screens if sufficient (i.e., biopsy-sized) additional donor tissue is available. The amount of tissue needed for a targeted rapid *ex vivo* screen is similar to the amount needed for an additional tumor biopsy, which facilitates translational applications. Other groups, e.g., Sato *et al.*, have successfully generated NEN organoid lines as excellent models for comprehensive mechanistic studies.^15^ However, the organoid expansion process took from several months to years^15^, limiting the option of feedback to the clinicians in the urgent scenario of rapidly progressive malignant GEP-NENs.

Our research lays the groundwork for prospective validation of patient-derived tumoroids as faithful *ex vivo* models for personalized screening of treatment efficacies. Larger prospective studies could evaluate the predictive relevance in more detail. Prospective studies could also more specifically test clinically applied cisplatin+etoposide and capecitabine+temozolomide combination therapy, which we did not include in our proof-of-concept study and could consider available second-line treatment options in the study protocol. Such studies can help identify more suitable second-line therapies in highly aggressive tumor types after first-line therapies fail. The standard processing workflow of tissues for PD tumoroid culture is compatible with multi-site sample collection – a prerequisite for prospective approaches in such rare malignancies.

Molecular drivers of the divergent clinical course of G3 NET and NEC are poorly understood, making individual treatment decisions challenging. Because advanced tumors often resist monotherapies, antineoplastic agents are combined to be more efficacious at lower doses.^29–33,37,39,40^ Patient-to-patient heterogeneity,^39^ intra-tumoral heterogeneity,^41^ and intracellular pathway dysregulation^42^ open new avenues for combining therapies to induce potent responses that monotherapy cannot achieve.^43^ We offer a strategy for using gene expression profiles to suggest treatments that can be combined with cisplatin chemotherapy; cisplatin-induced molecular stress response in high-grade GEP-NEN PD tumoroids is specific and mirrors perturbational effects (Fig. 4, Supplementary Fig. S5). We hypothesized that highly inverse signatures to cisplatin-monotherapy were likely to be ideal candidates for combinational treatments. Targeting these candidates and cisplatin may severely corrupt the cellular signaling state, killing cancer cells. We used these perturbational profiles to pinpoint two novel candidates for combinational therapy: Lysine Demethylase 5A (KDM5A) and interferon beta 1 (IFN1B) as (Fig. 4, Supplementary Fig. S5).

KDM5A is a histone demethylase that often represses target genes at transcriptional start sites^44^ and its role in neuroendocrine differentiation and tumorigenesis was recently described.^45,46^ Kaelin *et al.* demonstrated that Kdm5a promotes SCLC tumorigenesis *in vivo* and tumor proliferation and proposing inhibiting KDM5A as a therapeutic strategy.^46^ Genomic analysis of GEP-NENs has shown that in 45% - 52% of the tumors there is a KDM5A copy number gain.^10^ The findings of these two independent studies closely align with our finding that KDM5A plays a prominent role in neuroendocrine neoplasms. Upon combinational treatment of KDM5A inhibitor with cisplatin, three GEP-NENs we tested showed strong synergism and clinically attractive efficacies (Fig. 5, Supplementary Fig. S6).

Interestingly, KMD5A and cisplatin susceptibility have a functional relationship in lung adenocarcinoma, pointing towards altered chromatin regulation as a potential molecular mechanism for drug tolerance.^47^ Note that the sample in which the Cisplatin+KDM5A combination was ineffective had a mutational disfunction upstream of the H3K4 methylation axis. Mutations in lysine methyltransferase 2A (KMT2A) and menin (MEN1) regulate H3K4 methylation, so this dysfunction may have rendered the combination ineffective.

Type I interferons (IFN-α and IFN-β) are pro-inflammatory cytokines that can rapidly cause myriad downstream effects in tumor cells and promote antitumor immunity in immune cells.^48,49^ Type I interferons activate transcription factors of the signal transducer and activator of transcription (STAT) family, initiating protein synthesis from interferon-stimulated genes.^49^ Type 1 interferons are FDA-approved for mono- or combinational therapy because they cause tumor regression and may prolong survival in many other highly proliferative hematological and disseminated solid malignancies.^48^ IFN-α was used to treat advanced low-grade GEP-NETs^50–52^ but was superseded by other regimens (e.g., somatostatin analogs).^53^ Recently, two independent studies proposed that IFN-β be used to treat GEP-NETs because at low doses it effectively inhibits cell proliferation and stimulates apoptosis in cell lines *in vitro.^54,55^* In the clinically more relevant scenario of patient-derived high-grade GEP-NET tumoroids, we found IFNB1 was associated with the GEP-NEN perturbational signature. Exposure to Cisplatin+IFNB1 revealed they were synergistic and highly efficacious in treating a subset of highgrade GEP-NEN tumoroids (Fig. 5, Supplementary Fig. S6). This combinational approach may be an attractive option for patients with high-grade GEP-NETs, who now have few treatment options.^3,5^

Further studies in larger cohorts are needed to determine to what extent KDM5A- or IFNB1 combinations are NEC- or NET G3 specific and further efforts are needed to delineate the exact mechanisms behind treatment susceptibilities. Exact treatment schedules and/or therapeutic priming should also be evaluated *in vitro.* A recent extensive and comprehensive high-throughput combinational drug screen in breast, colon, and pancreatic cancer indicated that chemotherapeutics combined with apoptotic inducers or cell cycle inhibitors show promise for translational applications.^29^ Both KDM5A and IFNB1 fall into this category, and our study underlines their functional potency. KDM5A and IFNB1 may prove to be the Achilles Heels for high-grade GEP-NEN if combined with cisplatin.

In summary, we successfully cultured PD tumoroids of high-grade GEP-NENs for a rapid *ex vivo* drug screen. These tumoroids recapitulated key biological features of high-grade GEP-NEN and mimicked clinical response to cisplatin and temozolomide *ex vivo.* We also investigated molecular stress responses in PD tumoroids *in silico*, discovering and functionally validating Lysine demethylase 5A (KDM5A) and interferon-beta (IFNB1)—two vulnerabilities that act interact when combined with cisplatin. Either KDM5A or IFNB1 can be combined with cisplatin, opening new therapeutic options in high-grade GEP-NENs.

Our findings, that GEP-NEN PD tumoroids are promising candidates for rapid and biologically meaningful *ex vivo* pharmacotyping and that they can provide subsidiary therapy information, brings us closer to developing more personalized clinical protocols for later-line therapies in patients with aggressive high-grade GEP-NEN.

## Supporting information

SciScore

## ACKNOWLEDGMENTS

We thank the Tissue Bank Bern (Bern, Switzerland) and the Translational Research Unit (Institute of Pathology, Bern, Switzerland) for their technical, material, and administrative support and the Cytopathology (Institute of Pathology, Bern, Switzerland) for technical support performing formalin-fixation and paraffin embedding in tissue and cell culture material. We thank Dr. K. Tal for her editorial assistance. We thank Prof. S. Rottenberg (Institute of Animal Pathology, Vetsuisse Faculty, University of Bern, Switzerland) for his material support for the *ex vivo* and *in vivo* drug screens.

## AUTHOR CONTRIBUTIONS CRediT (ISSN 09531513)

**Conceptualization**: S.L.A.M., B.W., A.P.; **Methodology:** S.L.A.M., P.K.; **Software**: S.L.A.M., P.K., C.S.; **Validation**: S.L.A.M., K.D., P.K.; **Formal Analysis**: S.L.A.M., P.K.; **Investigation**: S.L.A.M., K.D., P.K., M.A.T., T.G., K.B., C.S., A. C.; **Resources**: S.L.A.M., A.K., C.A.K., D.H., J.S., M.C.S., I.M., B.W., A.P.; **Data Curation**: S.L.A.M., K.D., P.K., R.M.A.; **Writing—Original Draft Preparation**: S.L.A.M., K.D., B.W., A.P.; **Writing—Review and Editing**: S.L.A.M., K.D., B.W., A.P.; **Visualization**: S.L.A.M.; **Supervision**: S.L.A.M., B.W., A.P.; **Project Administration**: S.L.A.M., B. W., A.P.; **Funding Acquisition**: I.M., B.W., A.P.

All authors have read and agreed to the published version of the manuscript.

## SUPPLEMENTARY MATERIAL

**Supplementary Table 1:** Demographics and clinical data and microsatellite stability

**Supplementary Table 2:** Mutation profiling

**Supplementary Table 3:** Cytology of patient-derived tumoroids

**Supplementary Table 4:** Gene Ontology and GSEA of PD tumoroids vs original tumor tissue

**Supplementary Table 5:** GRmetrics

**Supplementary Table 6:** cMAP signatures

**Supplementary Figures S1 to S7**

**SUPPLEMENTARY FIGURE S1.**
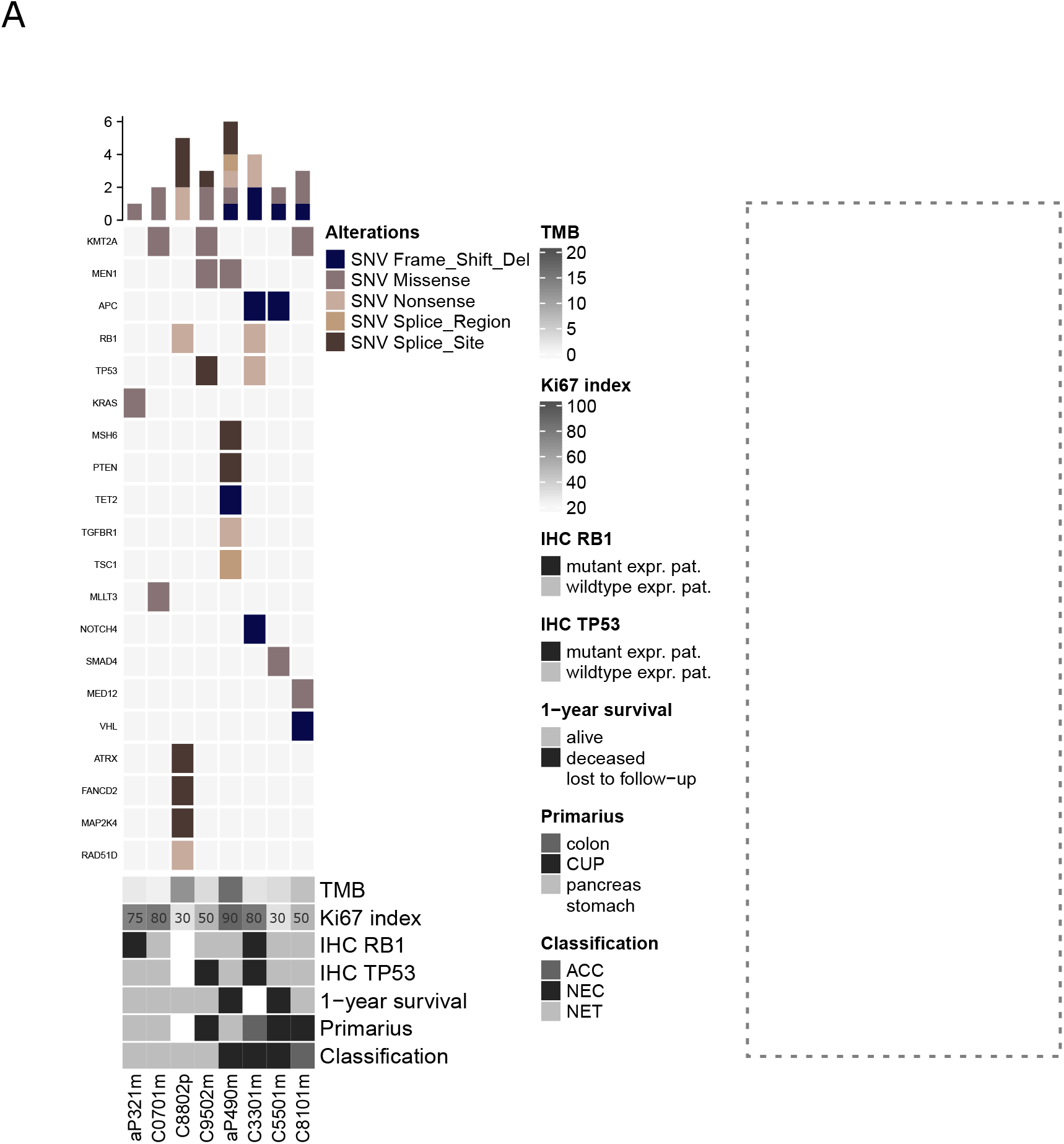
Detailed histomorphological-, mutational-, and clinical description of high-grade GEP-NEN patient cohort. **A)** Oncoplot showing all detected genes harboring alterations and a selection of clinicopathological parameters from GEP-NEN patients. The top panel indicates mutation counts (y-axis) per patient. The middle panel indicates types of single nucleotide variations (SNV) found in fresh frozen original tumor tissue from high-grade GEP-NEN patients. The lower panel indicates clinical parameters, including tumor mutational burden (TMB; mutation/Mb) and 1-year survival, IHC-based proliferation status (Ki-67; percent positive cells per tissue), RB1 protein expression, TP53 protein expression, location of primary tumor, and the diagnostic classification. NET neuroendocrine tumor, NEC neuroendocrine carcinoma, ACC acinar cell carcinoma, CUP cancer of unknown primary, MSI microsatellite instable, Mutant expr. pat. Mutant expression pattern (TP53 loss of protein (0% positive tumor cells) or overexpression (≥90% positive tumor cell; RB1 complete loss of protein), wildtype expr. pat. wildtype expression pattern.

**SUPPLEMENTARY FIGURE S2.**
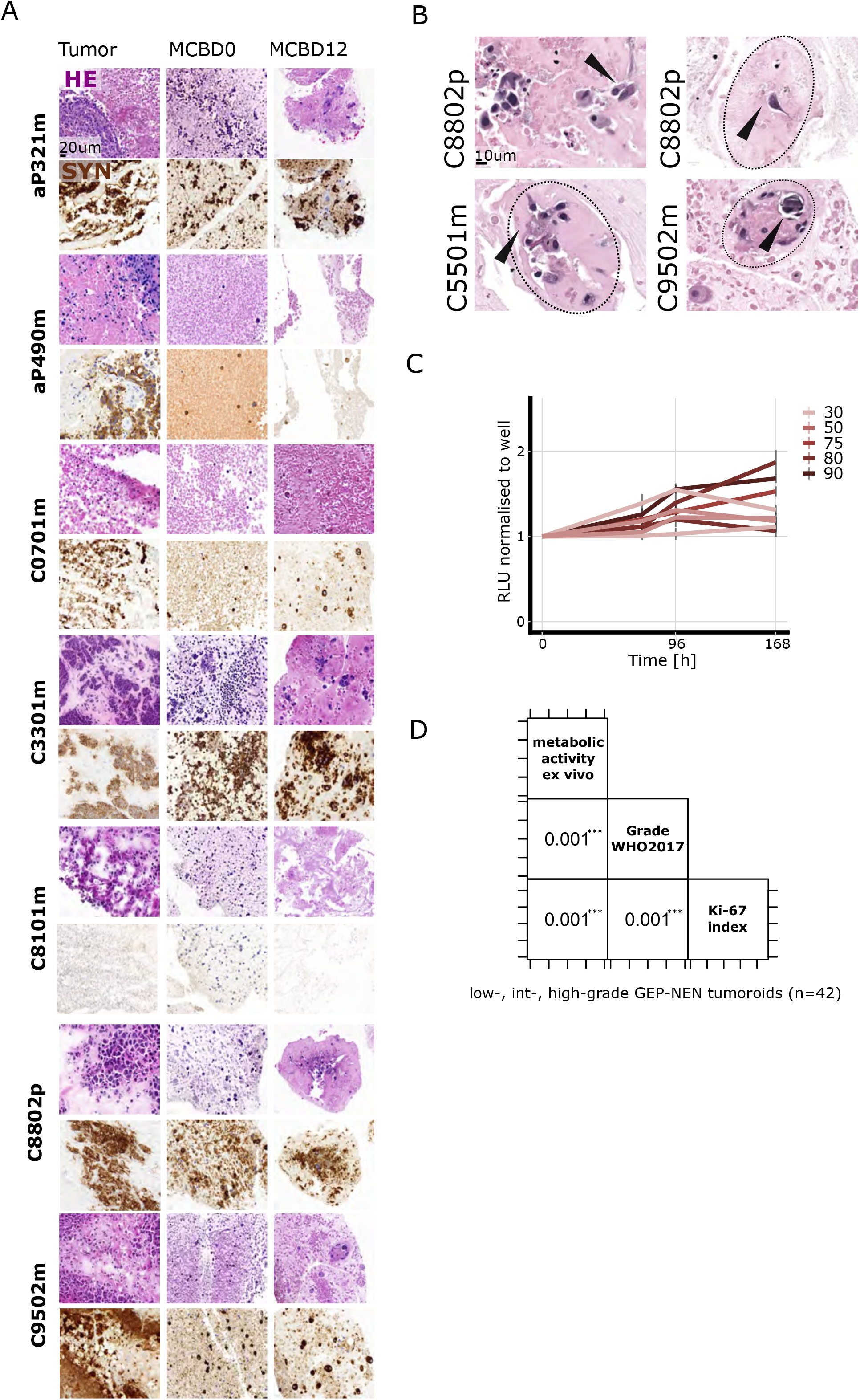
Histomorphological and functional characterization of patient-derived GEP-NEN tumoroids. **A)** Representative Hematoxylin and eosin (HE) stainings and synaptophysin (SYN) immunolabeling in original tumor tissue and patient-derived (PD) tumoroids. Cells were formalin-fixed and embedded into micro-cell-blocks directly after isolation (MCB D0) and after 12 days in culture (MCB D12). PD tumoroids and corresponding mirror blocks from fresh frozen original tumors were stained for HE or SYN. IHC slides were counterstained with hematoxylin. All stainings were assessed by two board-certified pathologists. Scale bar in all micrographs 20 μm. **B)** Representative HE staining in MCB of PD tumoroids after 12 days in culture. Black arrows and dotted lines highlight solitary fibroblasts (C8802p), focal deposition of extracellular matrix (C8802p, C5501m), or focal calcifications (C9502m). Scale bar in all micrographs 10 μm. **C)** Increase in metabolic activity measured by Real-time Glo (RTG) (a surrogate for cell proliferation/cell counts) in high-grade GEP-NEN patient-derived tumoroids. **D)** Correlation between metabolic activity (RTG) (a surrogate for cell proliferation/cell counts) and clinical parameters for proliferation (Grade; Ki-67 index) in low- (n=9), int- (n=23), and high-grade (n=10) GEP-NEN patient-derived tumoroids (number of patients n=42) (p-values are derived from χ2-test of independence using Monte Carlo simulation (n=1000)) RLU relative luminescence unit

**SUPPLEMENTARY FIGURE S3.**
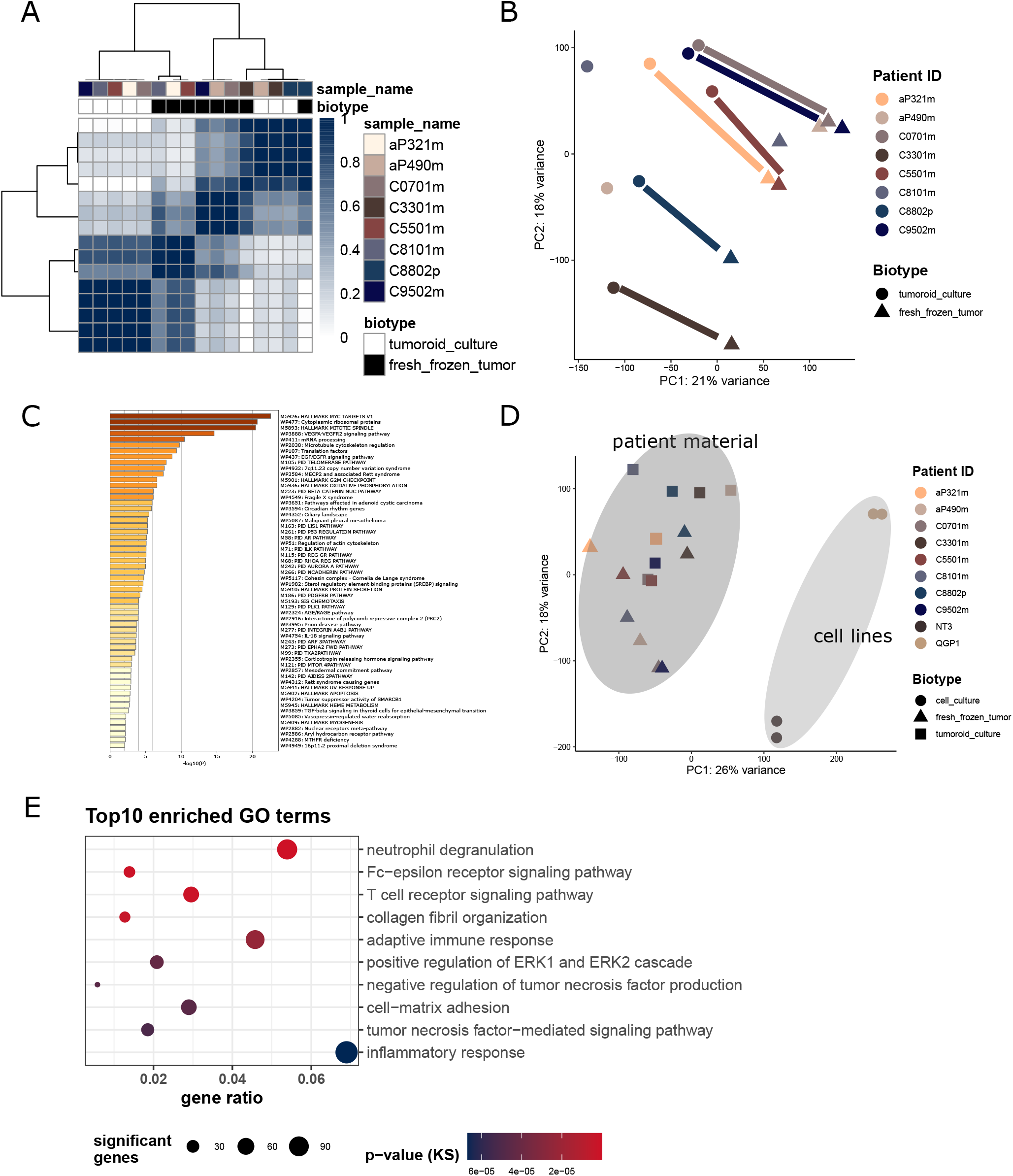
Transcriptional characterization of patient-derived GEP-NEN tumoroids. **A)** Consensus clustering on VIPER-inferred master regulator (MR) protein activity in patient-derived tumoroids and original tumor tissues. **B)** Principal component analysis (PCA) of gene expression in original tumor tissue and tissue-matched PD tumoroids. Patient-specific expression patterns are systematically retained in culture (solid lines). **C)** Statistically enriched pathways (Hallmark; Wiki; Canonical) in top 500 non-differentially expressed genes (Log2-fold change 1 ± 0.25) from control PD tumoroids and original tumor tissue. **D)** PCA of gene expression in original tumor tissue, tissue-matched PD tumoroids, and NEN cell line spheroids (NT3, QGP1). Gene expression in NEN cell line spheroids diverges from patient material and builds a separate cluster (light-grey circle). **E)** Top 10 significantly enriched gene ontology (GO) terms of biological processes (p-ks < 0.005). 8 out of 10 GO terms are related to the immune cell compartment. P-values were determined by the Kolmogorov-Smirnov test.

**SUPPLEMENTARY FIGURE S4.**
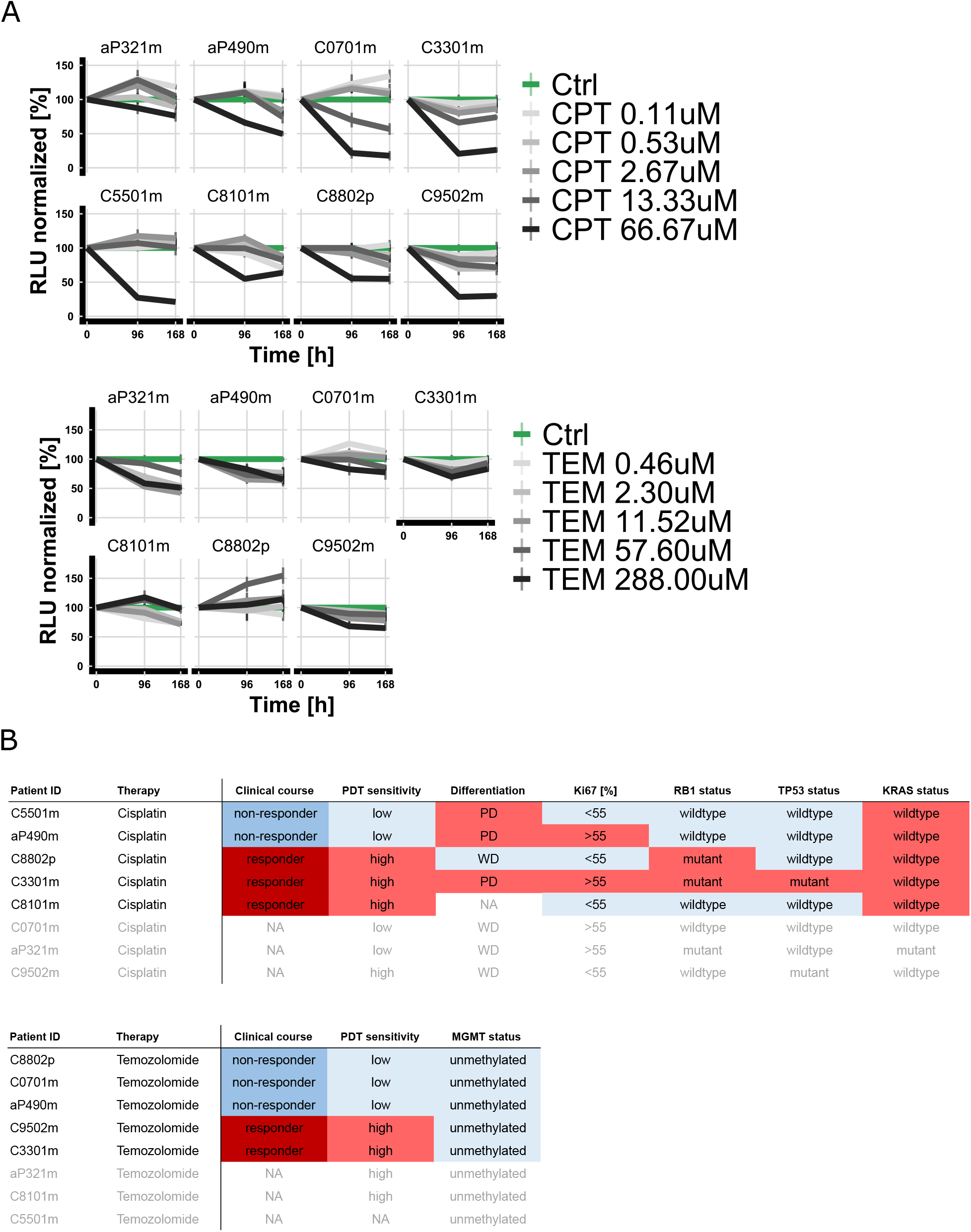
*ex vivo* pharmacotyping in patient-derived GEP-NEN tumoroids. **A)** *Ex vivo* viability curves from patient-derived (PD) tumoroids treated for 96 and 168 hours. Cisplatin (CPT) or temozolomide (TEM) data points are normalized to corresponding DMSO control (Ctrl) for each specific patient. Data represent mean ± SEM (n=1, three technical replicates). **B)** Overview of clinical course, *ex vivo* sensitivity in PD tumoroids, and clinicopathological features associated with clinical response.

**SUPPLEMENTARY FIGURE S5.**
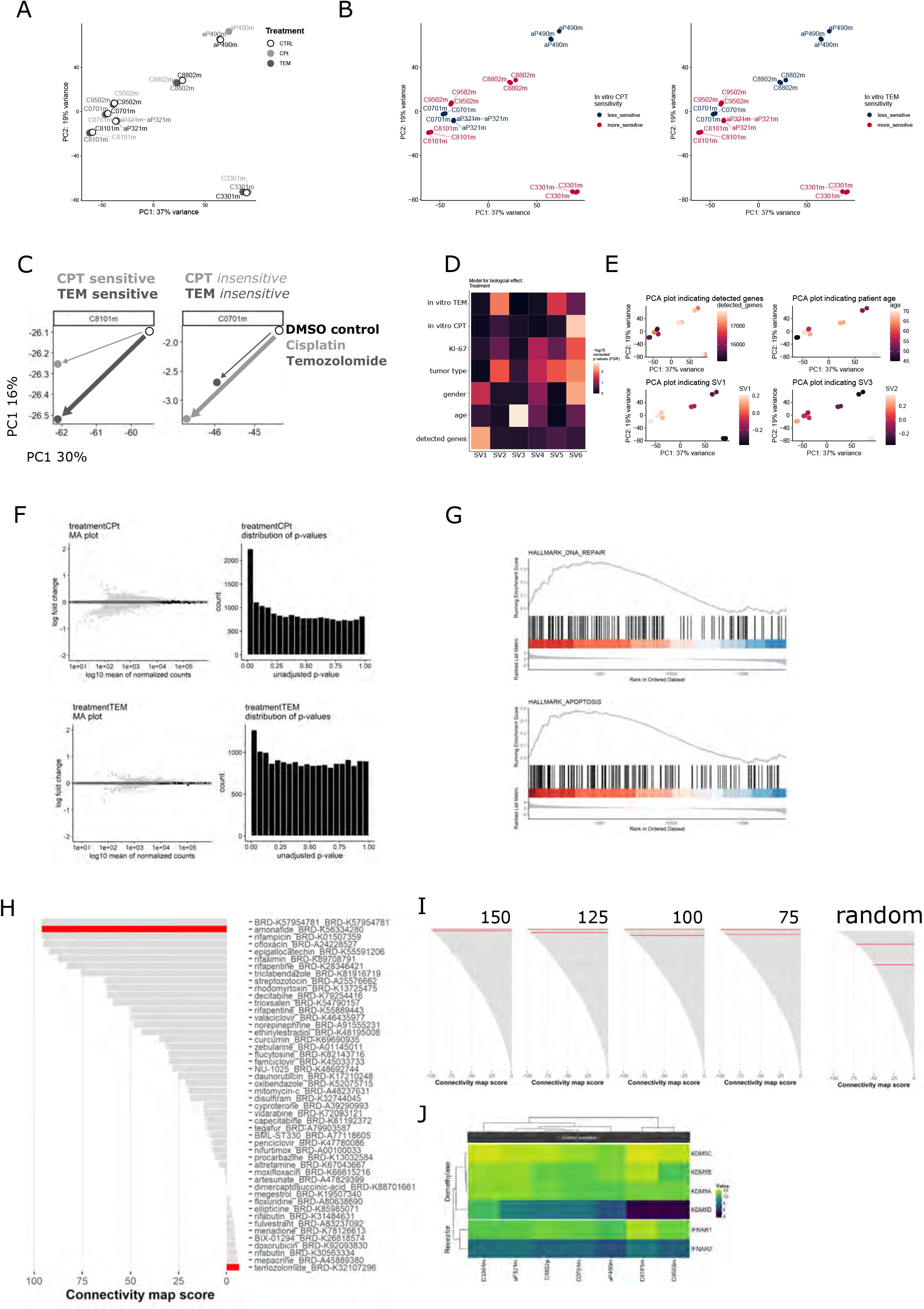
Characterization of molecular stress response in PD tumoroids. **A+B)** Principal component analysis (PCA) of gene expression in patient-derived (PD) tumoroids either treated with DMSO as a control (Ctrl) or treated with cisplatin (CPT) or temozolomide (TEM) at sublethal dosages. Parametrized *ex vivo* sensitivity is indicated by color. **C)** Representative PCA indicating the magnitude of gene expression change (top 2000 genes) in CPT or TEM-treated PD tumoroids either more sensitive (C8101m) or less sensitive (C0701m) for both treatments. **D)** Heatmap indicating association of known covariates with estimated surrogate variables (SV). Association was tested using linear models for continuous covariates and Kruskal-Wallis tests for categorical covariates. P-values were corrected for multiple testing using the false discovery rate (FDR) cutoff < 0.01. **E)** Representative PCA highlighting overlap of selected SVs with known covariates, including the number of detected genes and patient age. **F)** Mean-average (MA) plot of gene expression changes induced by treatment. Differential gene expression was compared between DMSO control PD tumoroids and PD tumoroids treated with sublethal dosages of CPT or TEM. Significantly differentially expressed genes (p-adj < 0.05) with an FDR < 0.1 are highlighted in light grey (left). A histogram of p-values for genes with mean-normalized-counts larger than 1 (right) indicates enrichment in significant p-values. **G)** Gene set enrichment plot showing enrichment of DNA repair and apoptosis in cisplatin-induced stress responses of PD tumoroids. The upper panel displays the running sum of gene sets and the leading edge. The lower panel represents log2 fold-change ranked detected genes in RNAseq. **H)** Waterfall plot of DNA-directed compounds in cMAP and their similarity (connectivity map score > 0) to the cisplatin-induced stress response. Amonafide (DNA intercalating agent) shows high similarity to cisplatin-induced stress response, whereas temozolomide (DNA alkylating agent) shows low similarity. **I)** Waterfall plot assessing the robustness of detecting KDM5A and IFNB1 in cMAP top inverse ranks. KDM5A and IFNB1 are highlighted in red. cMAP output remains stable upon systematic degradation of the number of input query genes. This ranking pattern was robust and specific: when input query signatures were systematically degraded (top 150/125/100/75 up- and downregulated DEGs (p-adj < 0.05), *IFNB1* and *KDM5A* remained among the top-ranked hits and had highly inverse connectivity map scores (τ < −91.00) while random permutation and selection of input genes to cMap database led to a complete loss of these ranks and insignificant connectivity scores (757/3195, τ = −49.47; 289/3195, τ = −71.76). **J)** Variance stabilizing transformation (VST)-normalized mRNA expression levels of KDM5 isoforms and IFNB1 receptor subunits (IFNAR1 and 2) in high-grade GEP-NEN patient-derived tumoroids.

**SUPPLEMENTARY FIGURE S6.**
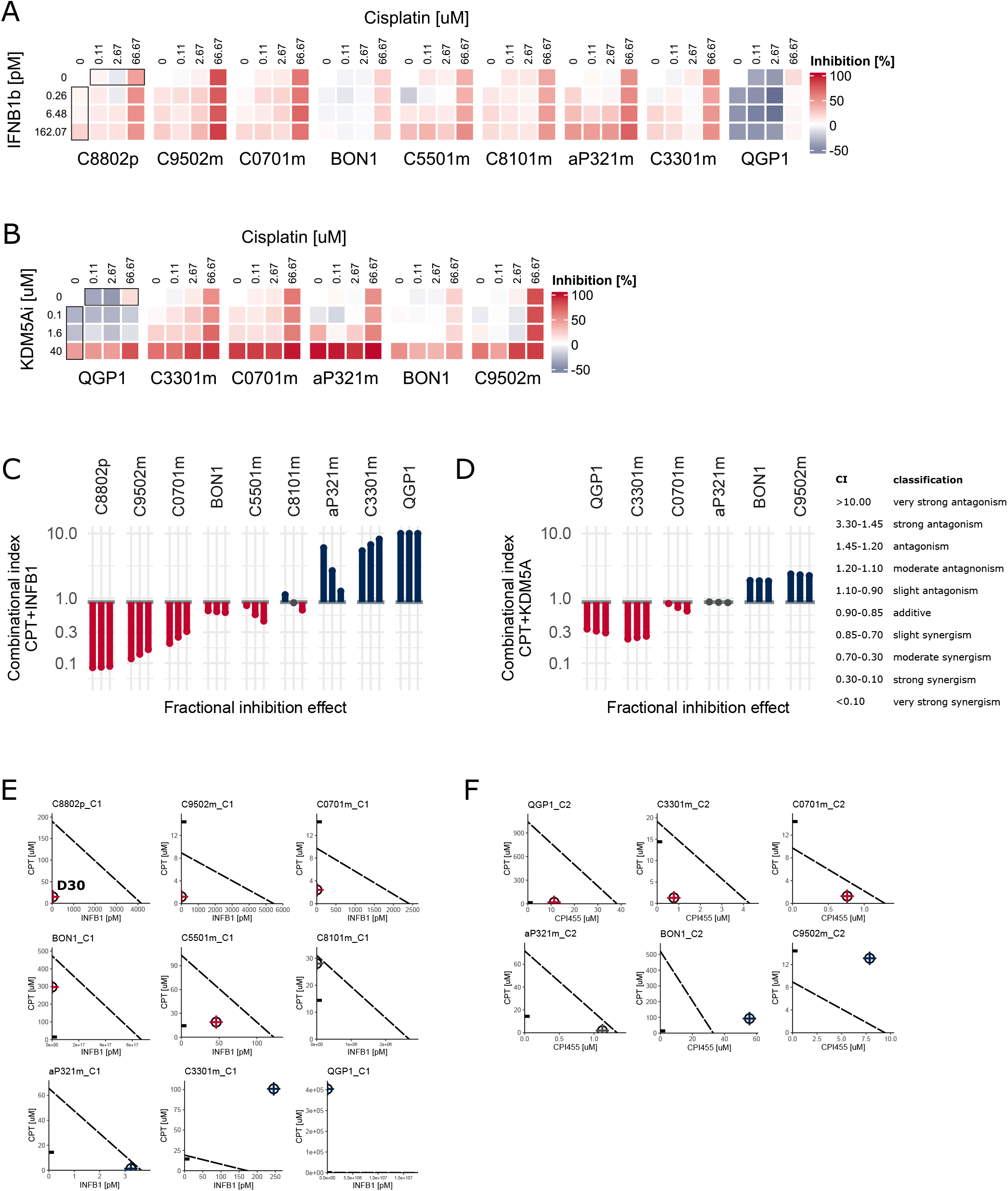
Assessment of combinational drug treatment in patient-derived tumoroids. **A+B)** Drug response heatmap of mono- and combinational treatment of cisplatin and recombinant IFNB1 or KDM5A inhibitor (CPI-455) in a full matrix design. The short-term treatment response was assessed after 24 hours, and inhibition was normalized to DMSO control for each sample. **C+D)** Bar graph indicating drug interaction between cisplatin and IFNB1 or KDM5A in PD tumoroids. Three representative fractional inhibition effect levels (0.25, 0.30, 0.35) were chosen based on the overall inhibitory effect of cisplatin monotherapy at physiologically relevant concentrations (14.4uM Cmax; inhibition 0.29). Shown are the combination index (CI) in red (synergistic), blue (antagonistic), or grey (zero-interaction). **E+F)** Isobologram showing drug interaction between cisplatin and IFNB1 or KDM5A in PD tumoroids at a 30% inhibition effect level. The dashed line indicates the zero-interaction isobole of isoactive monotherapies. Isoactive drug combinations are indicated by crossed circles in red (synergistic), blue (antagonistic), or grey (zero-interaction). The physiological dosage of cisplatin (cMax) is indicated by the black line on the y-axis.

**SUPPLEMENTARY FIGURE S7.**
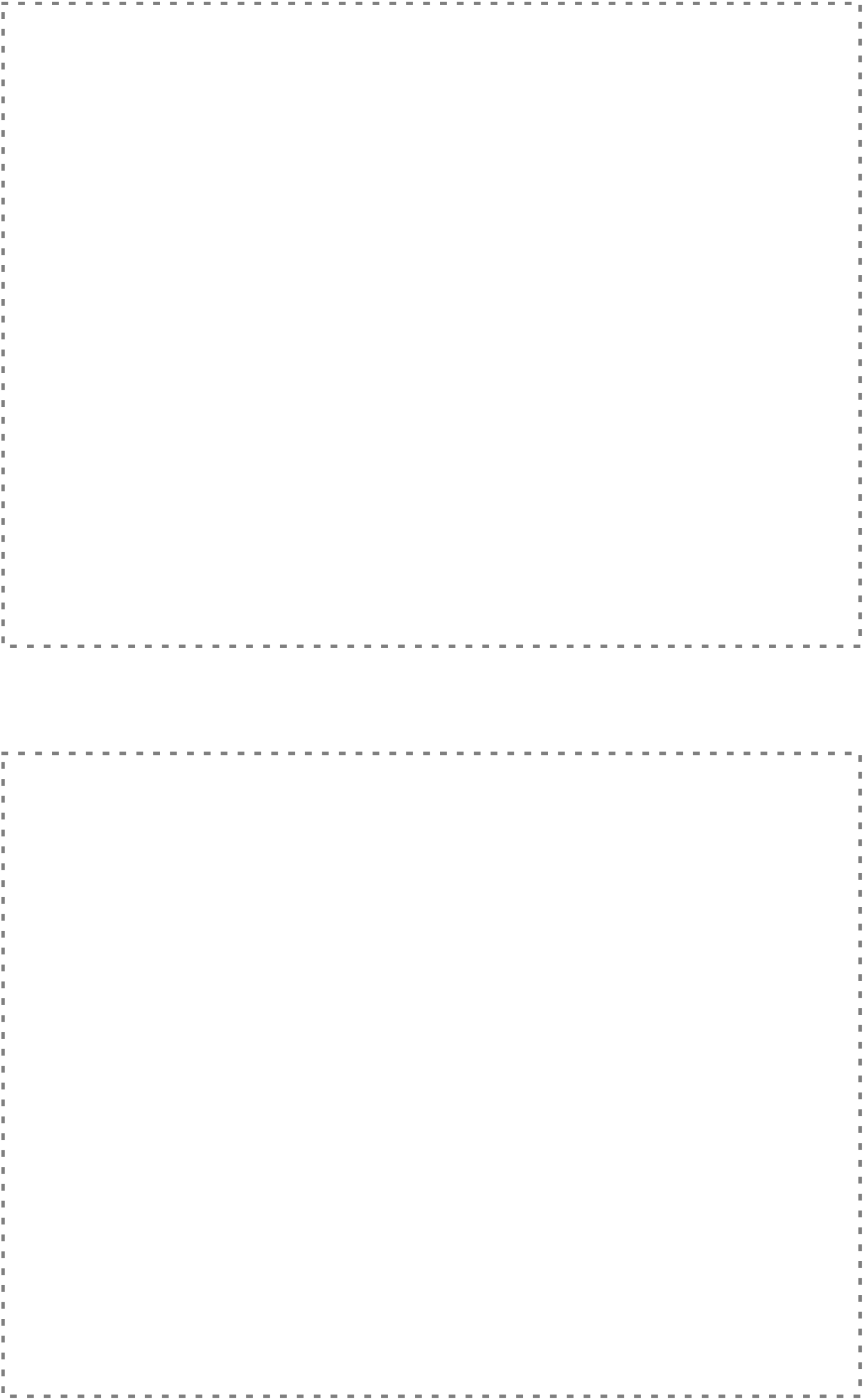
tbd

## Notes

**Financial support** The study was supported by the Swiss Cancer Research Foundation to A.P. (KFS-4227-08-2017) and Wilhelm Sander to I.M. and a generous donation by Dr. M. Gunzenhauser and B. Piepgras to B. W.

**Conflict of interest disclosure statement** The authors declare no potential conflicts of interest.

https://www.ncbi.nlm.nih.gov/geo/query/acc.cgi?acc=GSE98894

https://www.ncbi.nlm.nih.gov/geo/query/acc.cgi?acc=GSE213504

### Competing Interest Statement

The authors have declared no competing interest.

### Summary of Updates

Affiliations; Co-author list; SciScore; Improvment in figure size and figure legends and minor adaptions of preliminary data; Due to file size limitations a high resolution version of the manuscript and figures is available on request

## REFERENCES

1. Dasari, A., Mehta, K., Byers, L. A., Sorbye, H. & Yao, J. C. Comparative study of lung and extrapulmonary poorly differentiated neuroendocrine carcinomas: A SEER database analysis of 162,983 cases. Cancer 124, 807–815 (2018).

2. Dasari, A. et al. Trends in the Incidence, Prevalence, and Survival Outcomes in Patients With Neuroendocrine Tumors in the United States. JAMA Oncol. 3, 1335–1342 (2017).

3. Heetfeld, M. et al. Characteristics and treatment of patients with G3 gastroenteropancreatic neuroendocrine neoplasms. Endocr. Relat. Cancer 22, 657–664 (2015).

4. Sorbye, H. et al. Predictive and prognostic factors for treatment and survival in 305 patients with advanced gastrointestinal neuroendocrine carcinoma (WHO G3): the NORDIC NEC study. Ann. Oncol. Off. J. Eur. Soc. Med. Oncol. 24, 152–160 (2013).

5. Pavel, M. et al. Gastroenteropancreatic neuroendocrine neoplasms: ESMO Clinical Practice Guidelines for diagnosis, treatment and follow-up. Ann. Oncol. 31, 844–860 (2020).

6. Garcia-Carbonero, R. et al. ENETS Consensus Guidelines for High-Grade Gastroenteropancreatic Neuroendocrine Tumors and Neuroendocrine Carcinomas. Neuroendocrinology 103, 186–194 (2016).

7. Strosberg, J. R. et al. The NANETS Consensus Guidelines for the Diagnosis and Management of Poorly Differentiated (High-Grade) Extrapulmonary Neuroendocrine Carcinomas. Pancreas 39, 799–800 (2010).

8. Al-Toubah, T., Pelle, E., Valone, T., Haider, M. & Strosberg, J. R. Efficacy and Toxicity Analysis of Capecitabine and Temozolomide in Neuroendocrine Neoplasms. J. Natl. Compr. Canc. Netw. 20, 29–36 (2021).

9. Elvebakken, H. et al. A Consensus-Developed Morphological Re-Evaluation of 196 High-Grade Gastroenteropancreatic Neuroendocrine Neoplasms and Its Clinical Correlations. Neuroendocrinology 111, 883–894 (2021).

10. Venizelos, A. et al. The molecular characteristics of high-grade gastroenteropancreatic neuroendocrine neoplasms. Endocr. Relat. Cancer 29, 1–14 (2022).

11. Brennan, S. M. et al. Should extrapulmonary small cell cancer be managed like small cell lung cancer? Cancer 116, 888–895 (2010).

12. Rinke, A. et al. Treatment of advanced gastroenteropancreatic neuroendocrine neoplasia, are we on the way to personalised medicine? Gut 70, 1768–1781 (2021).

13. Detjen, K. et al. Models of Gastroenteropancreatic Neuroendocrine Neoplasms: Current Status and Future Directions. Neuroendocrinology 111, 217–236 (2021).

14. April-Monn, S. L. et al. Three-Dimensional Primary Cell Culture: A Novel Preclinical Model for Pancreatic Neuroendocrine Tumors. Neuroendocrinology 111, 273–287 (2021).

15. Kawasaki, K. et al. An Organoid Biobank of Neuroendocrine Neoplasms Enables Genotype-Phenotype Mapping. Cell 183, 1420–1435.e21 (2020).

16. Ben-David, U. et al. Patient-derived xenografts undergo mouse-specific tumor evolution. Nat. Genet. 49, 1567–1575 (2017).

17. April-Monn, S. L. et al. EZH2 Inhibition as New Epigenetic Treatment Option for Pancreatic Neuroendocrine Neoplasms (PanNENs). Cancers 13, 5014 (2021).

18. Liston, D. R. & Davis, M. Clinically Relevant Concentrations of Anticancer Drugs: A Guide for Nonclinical Studies. Clin. Cancer Res. 23, 3489–3498 (2017).

19. Chou, T.-C. Theoretical Basis, Experimental Design, and Computerized Simulation of Synergism and Antagonism in Drug Combination Studies. Pharmacol. Rev. 58, 621–681 (2006).

20. Driehuis, E., Kretzschmar, K. & Clevers, H. Establishment of patient-derived cancer organoids for drug-screening applications. Nat. Protoc. 1–30 (2020) doi:10.1038/s41596-020-0379-4.

21. Hafner, M., Niepel, M., Chung, M. & Sorger, P. K. Growth rate inhibition metrics correct for confounders in measuring sensitivity to cancer drugs. Nat. Methods 13, 521–527 (2016).

22. Chou, T.-C. Drug Combination Studies and Their Synergy Quantification Using the Chou-Talalay Method. Cancer Res. 70, 440–446 (2010).

23. Woo, J. H. et al. Elucidating Compound Mechanism of Action by Network Perturbation Analysis. Cell 162, 441–451 (2015).

24. Bansal, M. et al. A community computational challenge to predict the activity of pairs of compounds. Nat. Biotechnol. 32, 1213–1222 (2014).

25. Leek, J. T. & Storey, J. D. Capturing Heterogeneity in Gene Expression Studies by Surrogate Variable Analysis. PLOS Genet. 3, e161 (2007).

26. Alvarez, M. J. et al. A precision oncology approach to the pharmacological targeting of mechanistic dependencies in neuroendocrine tumors. Nat. Genet. (2018) doi:10.1038/s41588-018-0138-4.

27. Lamb, J. et al. The Connectivity Map: Using Gene-Expression Signatures to Connect Small Molecules, Genes, and Disease. Science 313, 1929–1935 (2006).

28. Kelland, L. The resurgence of platinum-based cancer chemotherapy. Nat. Rev. Cancer 7, 573–584 (2007).

29. Jaaks, P. et al. Effective drug combinations in breast, colon and pancreatic cancer cells. Nature 603, 166–173 (2022).

30. Sicklick, J. K. et al. Molecular profiling of cancer patients enables personalized combination therapy: the I-PREDICT study. Nat. Med. 25, 744–750 (2019).

31. Al-Lazikani, B., Banerji, U. & Workman, P. Combinatorial drug therapy for cancer in the post-genomic era. Nat. Biotechnol. 30, 679–692 (2012).

32. Ramaswamy, S. Rational Design of Cancer-Drug Combinations. N. Engl. J. Med. 357, 299–300 (2007).

33. Fitzgerald, J. B., Schoeberl, B., Nielsen, U. B. & Sorger, P. K. Systems biology and combination therapy in the quest for clinical efficacy. Nat. Chem. Biol. 2, 458–466 (2006).

34. Ooft, S. N. et al. Patient-derived organoids can predict response to chemotherapy in metastatic colorectal cancer patients. Sci. Transl. Med. 11, eaay2574 (2019).

35. Vlachogiannis, G. et al. Patient-derived organoids model treatment response of metastatic gastrointestinal cancers. Science 359, 920–926 (2018).

36. Tiriac, H. et al. Organoid Profiling Identifies Common Responders to Chemotherapy in Pancreatic Cancer. Cancer Discov. 8, 1112–1129 (2018).

37. Crystal, A. S. et al. Patient-derived models of acquired resistance can identify effective drug combinations for cancer. Science 346, 1480–1486 (2014).

38. Zehir, A. et al. Mutational landscape of metastatic cancer revealed from prospective clinical sequencing of 10,000 patients. Nat. Med. 23, 703–713 (2017).

39. Palmer, A. C. & Sorger, P. K. Combination Cancer Therapy Can Confer Benefit via Patient-to-Patient Variability without Drug Additivity or Synergy. Cell 171, 1678–1691.e13 (2017).

40. Gillies, R. J., Verduzco, D. & Gatenby, R. A. Evolutionary dynamics of carcinogenesis and why targeted therapy does not work. Nat. Rev. Cancer 12, 487–493 (2012).

41. Gerlinger, M. et al. Intratumor Heterogeneity and Branched Evolution Revealed by Multiregion Sequencing. N. Engl. J. Med. 366, 883–892 (2012).

42. Hyman, D. M., Taylor, B. S. & Baselga, J. Implementing Genome-Driven Oncology. Cell 168, 584–599 (2017).

43. Diaz, J. E. et al. The transcriptomic response of cells to a drug combination is more than the sum of the responses to the monotherapies. eLife 9, e52707 (2020).

44. Pedersen, M. T. & Helin, K. Histone demethylases in development and disease. Trends Cell Biol. 20, 662–671 (2010).

45. Oser, M. G. et al. The KDM5A/RBP2 histone demethylase represses NOTCH signaling to sustain neuroendocrine differentiation and promote small cell lung cancer tumorigenesis. Genes Dev. 33, 1718–1738 (2019).

46. Lin, W. et al. Loss of the retinoblastoma binding protein 2 (RBP2) histone demethylase suppresses tumorigenesis in mice lacking Rb1 or Men1. Proc. Natl. Acad. Sci. 108, 13379–13386 (2011).

47. Sharma, S. V. et al. A Chromatin-Mediated Reversible Drug-Tolerant State in Cancer Cell Subpopulations. Cell 141, 69–80 (2010).

48. Borden, E. C. et al. Interferons at age 50: past, current and future impact on biomedicine. Nat. Rev. Drug Discov. 6, 975–990 (2007).

49. Stark, G. R. & Darnell, J. E. The JAK-STAT Pathway at Twenty. Immunity 36, 503–514 (2012).

50. Eriksson, B. & Oberg, K. An update of the medical treatment of malignant endocrine pancreatic tumors. Acta Oncol. 32, 203–208 (1993).

51. Oberg, K. & Eriksson, B. The role of interferons in the management of carcinoid tumours. Br. J. Haematol. 79, 74–77 (1991).

52. Creutzfeldt, W., Bartsch, H. H., Jacubaschke, U. & Stöckmann, F. Treatment of gastrointestinal endocrine tumours with interferon-alpha and octreotide. Acta Oncol. 30, 529–535 (1991).

53. Kvols, L. K. et al. Treatment of the malignant carcinoid syndrome. Evaluation of a long-acting somatostatin analogue. N. Engl. J. Med. 315, 663–666 (1986).

54. Vitale, G. et al. IFN-β Is a Highly Potent Inhibitor of Gastroenteropancreatic Neuroendocrine Tumor Cell Growth In vitro. Cancer Res. 66, 554–562 (2006).

55. Zitzmann, K. et al. SOCS1 Silencing Enhances Antitumor Activity of Type I IFNs by Regulating Apoptosis in Neuroendocrine Tumor Cells. Cancer Res. 67, 5025–5032 (2007).

