## Supplementary material for "Patient-derived tumoroids of advanced high-grade neuroendocrine neoplasms mimic patient chemotherapy responses and guide the design of personalized combination therapies": SciScore

SciScore: 9
What's this?

Document Identifier: **106181**

### SciScore Report

Below you will find your SciScore report containing three tables. Your score is calculated based on adherence to scientific rigor criteria (Table 1) and identification of key biological resources (Table 2). Table 3 contains statistical tests and oligonucleotides but is not scored. If SciScore makes any mistakes, please contact us to help us learn and improve.

**Table 1: Rigor Adherence Table**

| Ethics |
|---|
| Field Sample Permit: The specimens were processed as described in April-Monn , et al ( 2021). |
| IRB: KEK-BE 105/2015 ) in accord with the Swiss Federal Human Research Act and by the ethics committee at Charité Universitätsmedizin Berlin ( Ref . -Nr . |
| Consent: All patients included in the study signed an institutional informed consent and agreed on the use of residual material. |
| Inclusion and Exclusion Criteria |
| Inclusion criteria were histopathologic diagnosis of G3 gastroenteropancreatic neuroendocrine neoplasm , availability of both tumor tissue- and matching cryomaterial for ex vivo culture , and tumor purity of >70 % . |
| Attrition |
| not detected. |
| Sex as a biological variable |
| The cohort included 3 female and 5 male patients; their ages varied from 39 to 70 years ( mean = 58.0; SD = 11.8) . |
| Subject Demographics |
| Age: The cohort included 3 female and 5 male patients; their ages varied from 39 to 70 years ( mean = 58.0; SD = 11.8) . |
| Randomization |
| not detected. |
| Blinding |
| A blinded experimenter scored ex vivo experiments and sensitivities to treatments . |
| Power Analysis |
| not detected. |
| Replication |

| | |
|---|---|
| Identifiers: Nucleic acid quantification was performed with the Qubit DNA/RNA HS detection kit ( Thermo Fisher Scientific , #Q32852) . | #Q32852 |
| Identifiers: 26 We leveraged a pan-GEP-NEN regulatory network ( context-specific interactome ) from transcriptional profiles of GEP-NEN patient samples ( n=212 ) ( GSE98894) . | GSE98894 |
| Identifiers: Data availability Sequence data that support the findings of this study have been deposited in Gene Expression Omnibus (GEO); primary accession code is GSE213504. | GSE213504 |

**Table 2: Key Resources Table**

| Your Sentences | REAGENT or RESOURCE | SOURCE | IDENTIFIER |
|---|---|---|---|
| Antibodies | | | |
| Visualization used a Bond Polymer Refine Detection Kit ( Leica , #DS9800 ) ( RRID:AB_2891238 ) for visualization; DAB ( 3,3#-Diaminobenzidine ) was the chromogen . | Bond Polymer Refine Detection | Leica Biosystems | (Leica Biosystems Cat# DS9800, RRID:AB_2891238)(link ) |
| Experimental Models: Cell Lines | | | |
| NEN cell line culture: The QGP1 cell line ( RRID:CVCL_3143 ) was purchased from the Japanese Health Sciences Foundation in 2011 . | QGP1 | | ( RRID:CVCL_3143)(link ) |
| The BON1 cell line ( RRID:CVCL_3985 ) was provided by E.J . M. Speel in 2011 . | The BON1 cell | | ( RRID:CVCL_3985)(link ) |
| QGP1 cells were authenticated by their specific cancer cell profile. | QGP1 | | Suggestion: JCRB Cat# JCRB0183, RRID:CVCL_3143( link) |
| Software and Algorithms | | | |
| Sources of population frequencies that were used for auto-classification of benign variation include gnomAD ( RRID:SCR_014964 ) and ExAC ( RRID:SCR_004068) . | gnomAD | | Genome Aggregation Database ( RRID:SCR_014964)(link ) |
| | ExAC | | ExAc ( RRID:SCR_004068)(link ) |
| We retrieved annotations of oncogenic effects of identified variants from the OncoKB precision oncology knowledge database ( RRID:SCR_014782 ) and assessed known activating mutations in oncogenes and inactivating mutations in tumor suppressors ( Tier 1 and Tier 2) . | the OncoKB | | OncoKB ( RRID:SCR_014782)(link ) |
| OncoPrint function from ComplexHeatmap v2.6.2 ( PMID 27207943 ) ( RRID:SCR_017270 ) was used for visualization . | ComplexHeatmap | | ComplexHeatmap ( RRID:SCR_017270)(link ) |
| Reads were demultiplexed and converted to FASTQ format with bcl2fastq v2.20.0.422 ( RRID:SCR_015058) . | FASTQ | | bcl2fastq ( RRID:SCR_015058)(link ) |
| Cutadapt v2.5 ( PMID 28715235 ) ( RRID:SCR_011841 ) was used to trim Illumina adapter sequences and mask 3' homopolymers longer than 10 bp . | Cutadapt | | cutadapt ( RRID:SCR_011841)(link ) |
| RRID:SCR_010910); mapping reads to this custom list were discarded. | RRID:SCR_010910); | | BWA ( RRID:SCR_010910)(link ) |

| | | | |
|---|---|---|---|
| At each step , we used FastQC v0.11.7 ( RRID:SCR_014583 ) to track read quality . | FastQC | | FastQC ( RRID:SCR_014583)(link ) |
| Processed reads were mapped to the human genome ( GRCh37 , GENCODE annotation v37 ) with STAR v2.7.3a ( PMID 23104886 ) ( RRID:SCR_004463) | STAR | | STAR ( RRID:SCR_004463)(link ) |
| Mapped reads were deduplicated based on the 8bp UMI in the R2; we used UMI-tools v0.5 ( PMID 28100584 ) ( RRID:SCR_017048 ) and the default directional method. | UMI-tools | | UMI-tools ( RRID:SCR_017048)(link ) |
| Deduplicated reads were assigned to GENCODE v37 genes in subread v2.0.1 ( PMID 24227677 ) ( RRID:SCR_009803) . | in subread | | Subread ( RRID:SCR_009803)(link ) |
| To determine differentially expressed genes , we combined limma voom ( PMID 25605792 ) ( RRID:SCR_010943 ) with the duplicateCorrelation function to model repeated measurements of the same patient . | limma | | LIMMA ( RRID:SCR_010943)(link ) |
| For drug-treated tumoroids , we determined differential expression with DESeq2 v1.32.0 ( PMID 25516281 ) ( RRID:SCR_000154) . | DESeq2 | | DESeq ( RRID:SCR_000154)(link ) |
| Treatment-independent expression variability was modeled using surrogate variable analysis ( SVA ) from sva v3.40.0 25 ( PMID 17907809 ) ( RRID:SCR_002155) . | SVA sva | | SVA ( RRID:SCR_002155)(link ) |
| All available surrogate variables were added to the DESeq2 model . | DESeq2 | | Suggestion: (DESeq, RRID:SCR_000154)( link) |
| Original tumor tissues and tumoroids were consensus clustered with ConsensusClusterPlus v1.54.0 ( PMID 20427518 ) ( RRID:SCR_016954 ) on the Pearson correlation of the 2000 most variable genes ( innerLinkage: Ward.D2 , finalLinkage: Average ) . | consensus | | ConsensusClusterPlus ( RRID:SCR_016954)(link ) |
| We used ARACNe ( accurate reconstruction of cellular networks ) ( PMID 15778709 ) ( RRID:SCR_002180 ) to reverse-engineer regulatory networks and VIPER ( Virtual Proteomic by Enriched Regulon analysis)26 to transform transcriptional profiles into master regulators protein activity profiles and to infer master regulator protein activity in original tumors and patient-derived tumoroids ( n=8 ) from our GEP-NEN patients . | ARACNe | | ARACNE ( RRID:SCR_002180)(link ) |

| | | | |
|---|---|---|---|
| To compare original tumor tissue and tumoroids, we selected differentially expressed genes (adjusted p-value < 0.05), and tested enrichment of Gene Ontology terms in topGo v2.44.0 (PMID: 16606683) (RRID:SCR_014798) (Kolmogorov-Smirnov, adjusted p-value < 0.01). | topGo | | topGO ( RRID:SCR_014798)(link ) |
| Gene set enrichment analysis (GSEA) (RRID:SCR_003199) was performed in clusterProfiler v. | Gene set enrichment analysis | | Gene Set Enrichment Analysis ( RRID:SCR_003199)(link ) |
| 3.18.1 (PMID: 22455463) (RRID:SCR_016884) based on log2 expression fold changes. | | | clusterProfiler ( RRID:SCR_016884)(link ) |
| Perturbational profiling in cMap: We compared the top and bottom 150 genes from drug versus control-treated tumoroids (adjusted p-value < 0.05, sorted by the Wald statistic) to the compendium of perturbational reference signatures from Connectivity Map (L1000, Touchstone v1.0)27 (RRID:SCR_015674) and extracted connectivity map scores (τ) for all available knock-down (kd), overexpression (oe), and compound perturbagens. | cMap: | | Connectivity Map 02 ( RRID:SCR_015674)(link ) |
| Data availability Sequence data that support the findings of this study have been deposited in Gene Expression Omnibus (GEO); primary accession code is GSE213504. | Gene Expression Omnibus | | Suggestion: (Gene Expression Omnibus (GEO), RRID:SCR_005012)( link) |

**Other Entities Detected**

| Your Sentences | Recognized Entity |
|---|---|
| Statistical Tests | |
| Original tumor tissues and tumoroids were consensus clustered with ConsensusClusterPlus v1.54.0 ( PMID 20427518 ) ( RRID:SCR_016954 ) on the Pearson correlation of the 2000 most variable genes ( innerLinkage: Ward.D2 , finalLinkage: Average ) . | Pearson correlation |
| To compare original tumor tissue and tumoroids, we selected differentially expressed genes (adjusted p-value < 0.05), and tested enrichment of Gene Ontology terms in topGo v2.44.0 (PMID: 16606683) (RRID:SCR_014798) (Kolmogorov-Smirnov, adjusted p-value < 0.01). | Kolmogorov-Smirnov |

SciScore is an automated tool that is designed to assist expert reviewers by finding and presenting formulaic information scattered throughout a paper in a standard, easy to digest format. ***SciScore is not a substitute for expert review.*** SciScore also checks for the presence and correctness of several unique identifiers, including RRIDs (research resource identifiers) in the manuscript, detects sentences that appear to be missing RRIDs, and can even suggest RRIDs under certain circumstances. **All RRID suggestions should be verified;** only the author can know whether the suggestions are correct.

For a full description of scored criteria and tips for improving your score, please see https://www.scicrunch.com/sciscorereport-faq
